# Hierarchical design of multi-scale protein complexes by combinatorial assembly of oligomeric helical bundle and repeat protein building blocks

**DOI:** 10.1101/2020.07.27.221333

**Authors:** Yang Hsia, Rubul Mout, William Sheffler, Natasha I. Edman, Ivan Vulovic, Young-Jun Park, Rachel L. Redler, Matthew J. Bick, Asim K. Bera, Alexis Courbet, Alex Kang, TJ Brunette, Una Nattermann, Evelyn Tsai, Ayesha Saleem, Cameron M. Chow, Damian Ekiert, Gira Bhabha, David Veesler, David Baker

## Abstract

A goal of *de novo* protein design is to develop a systematic and robust approach to generating complex nanomaterials from stable building blocks. Due to their structural regularity and simplicity, a wide range of monomeric repeat proteins and oligomeric helical bundle structures have been designed and characterized. Here we describe a stepwise hierarchical approach to building up multi-component symmetric protein assemblies using these structures. We first connect designed helical repeat proteins (DHRs) to designed helical bundle proteins (HBs) to generate a large library of heterodimeric and homooligomeric building blocks; the latter have cyclic symmetries ranging from C2 to C6. All of the building blocks have repeat proteins with accessible termini, which we take advantage of in a second round of architecture guided rigid helical fusion (WORMS) to generate larger symmetric assemblies including C3 and C5 cyclic and D2 dihedral rings, a tetrahedral cage, and a 120 subunit icosahedral cage. Characterization of the structures by small angle x-ray scattering, x-ray crystallography, and cryo-electron microscopy demonstrates that the hierarchical design approach can accurately and robustly generate a wide range of macromolecular assemblies; with a diameter of 43nm, the icosahedral nanocage is the largest structurally validated designed cage to date. The computational methods and building block sets described here provide a very general route to new *de novo* designed symmetric protein nanomaterials.

## Introduction

There has been considerable recent interest in designing self assembling protein nano structures and materials^1,2^. Computational protein design has been used to create proteins that self-assemble into a wide variety of higher order structures, from cyclic^3^ and dihedral symmetries^4^ to point group nanocages^5–7^, 1-dimensional fibers^8^, and 2-dimensional arrays^9^. The nanocages have been utilized in vaccine development^10,11^, drug delivery^12^, and as microscopy standards^7^. Most of these structures have been created by symmetrically docking protein building blocks followed by sequence optimization at the new interfaces^3,5–7,9,13^ using RosettaDesign^14^. However, interface design remains challenging, and designable interface quality is heavily dependent on how well the building blocks complement each other during design. An alternative approach which avoids the need for designing new interfaces is to fuse oligomeric protein building blocks with helical linkers; while this has led to a number of new materials^15^, lack of rigidity has made the structures of these assemblies difficult to precisely specify. More rigid junctions created by overlapping ideal helices and designing around the junction region has resulted in more predictable structures^16,17^, including closed ring dihedral structures which require even more precise structure predictions^18^. This rigid fusion method, however, has its own set of challenges in comparison to designing a new non-covalent protein-protein interface: first, for any pair of protein building blocks, there are far fewer positions for rigid fusion than are for unconstrained protein-protein docking limiting the space of possible solutions, and second, while in the non-covalent protein interface case the space searched can be limited by restricting building blocks to the symmetry axes of the desired nanomaterial, this is not possible in the case of rigid fusions, making the search more difficult as the number of building blocks increases.

A potential solution to the issue of having smaller numbers of possible fusion positions for a given pair of building blocks in the rigid helix fusion method is to systematically generate large numbers of building blocks having properties ideal for helix fusion. Attractive candidates for such an approach are *de novo* helical repeat proteins (DHRs)^22^ consisting of a tandemly repeated structural unit, which provide a wide range of struts of different shape and curvature for building nanomaterials, and parametric helical bundles (HBs)^19–22^ which provide a wide range of preformed protein-protein interfaces for locking together different protein subunits in a designed nanomaterial. Many examples of both classes of designed proteins have been solved by x-ray crystallography, and they are typically very stable. We reasoned that by systematically fusing DHR “arms” to central HB “hubs” we could generate building blocks with a wide range of geometries and valencies that, because of the modular nature of repeat proteins, enable a very large number of rigid helix fusions: given two such building blocks with N- and C-terminally extending repeat protein arms, the potentially rigid fusion sites are any pair of internal helical residues in the DHR arms.

With a large library of building blocks, the challenge is then to develop a method to very quickly traverse all possible combinations of fusion locations. We present here WORMS, a software package that uses geometric hashing of transforms to very quickly and systematically identify the fusion positions in large sets of building blocks that generate any specified symmetric architecture, and describe the use of the software to design a broad range of symmetric assemblies.

## Results

We describe the development of methods for creating large and modular libraries of building blocks by fusing DHRs to HBs, and then using them to generate symmetric assemblies by rapidly scanning through the combinatorially large number of possible rigid helix fusions for those generating the desired architecture. We present the new methodology and results in two sections. In section one, we describe the systematic generation of homo- and hetero-oligomeric building blocks from *de novo* designed helical bundles, helical oligomers, and repeat proteins (Figure 1a). In the second section, we describe the use of these building blocks to assemble a wide variety of higher order symmetric architectures (Figure 1c).

**Figure 1.**
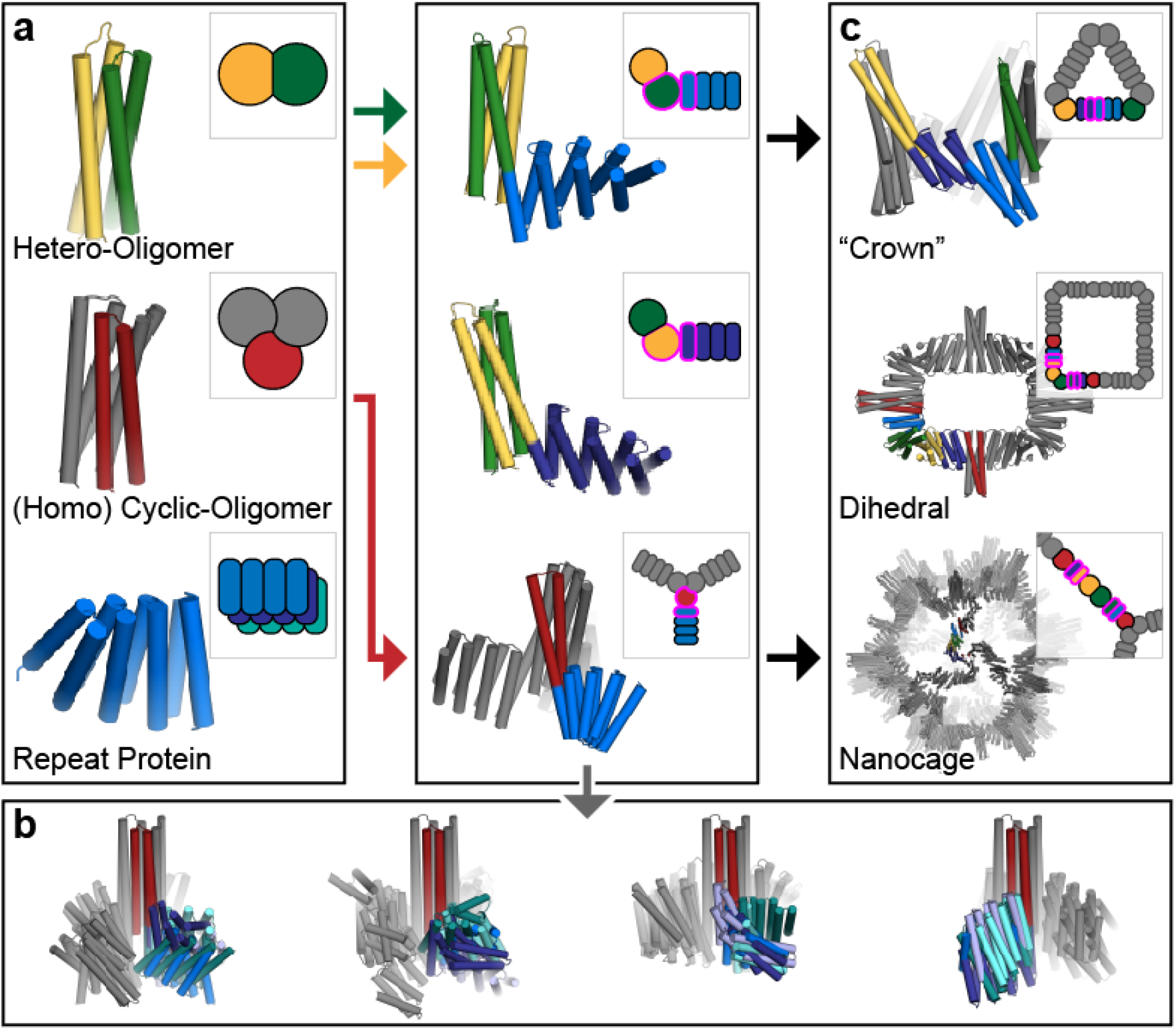
Overview of the rigid hierarchical fusion approach. (a) Hetero- (yellow/green) and homo- (red) oligomeric helical bundles are fused to *de novo* helical repeat proteins (shades of blue) (left) to create a wide range of building blocks using HelixDock and HelixFuse (center). Symmetric units shown in grey. (b) Twenty representative HelixFuse outputs overlaid in groups of five to display the wide range of diversity that can be generated by using a single helical bundle core. (c) These are then further assembled into higher ordered structures through helical fusion (WORMS, right). The examples shown are cyclic crowns (top), dihedral rings (middle), and icosahedral nanocages (bottom).

### Section 1: Systematic generation of oligomeric building blocks

To generate a wide variety of building blocks, we explored two different methodologies for fusing DHRs to HBs (Figure 1a). The first is to dock the DHR units to the HBs, redesign the residues at the newly created interface, and then build loops between nearby termini (HelixDock, HD). The second protocol simplifies the process by overlapping the helical termini of the DHRs and HBs and designing only the immediate residues around the junction (HelixFuse, HF). As an example of the combinatorial diversity that can be generated due to the large number of possible internal helical fusion sites in a DHR (nearly all helical residues), a single terminus from a single helical bundle (2L6HC3-12^20^, N-terminus) combined with the library of 44 verified DHRs resulted in 259 different structures (Figure 1b).

#### HelixDock (HD) approach

44 DHRs with validated structures^23^ and 11 HBs^20,24^ (including some without pre-verified structures) were selected as input scaffolds for symmetrical docking using a modified version of the sicdock software^3^. In each case, N copies of the DHR, one for each monomer in the helical bundle, were symmetrically docked onto the HB, sampling all six degrees of freedom, to generate star shaped structures with repeat protein arms emanating symmetrically from the helical bundle in the center. Docked configurations with linkable N- and C-termini within a distance cutoff of 9Å with interfaces predicted to yield low energy designs^25^ were then subjected to Rosetta sequence design to optimize the residue identity and packing at the newly formed interface. Designs with high predicted domain-domain binding energy and shape complementarity^26^ were identified, and loops connecting chain the termini were built using the ConnectChainsMover^17^. Structures with good loop geometry (passing worse9merFilter and FoldabilityFilter) were forward folded with RosettaRemodel^27^ symmetrically, and those with sequences which fold into the designed structure *in silico* were identified.

Synthetic genes encoding a subset of the selected designs with a wide range of shapes were synthesized and the proteins expressed in *E. coli*. Of the 115 sequences ordered successfully synthesized, 65 resulted in soluble protein. Those with poor expression and/or solution behavior were discarded. Of the remaining, 39 had relatively monodisperse Size Exclusion Chromatography (SEC) profiles that matched what was expected from the design. Of the ones selected for small angle X-ray scattering (SAXS), 17 had profiles close to those computed from the design models (Figure S1-3). Design C3_HD-1069, was crystallized and solved to 2.4 Å (Figure 2a). Although the two loops connecting to the HB are unresolved in the structure, the resulting placement of the DHR remains correct (unresolved loops were also present in the original HB structure (2L6HC3_6)^20^. The resolved rotamers at the newly designed interface between the HB and DHR are also as designed.

**Figure 2.**
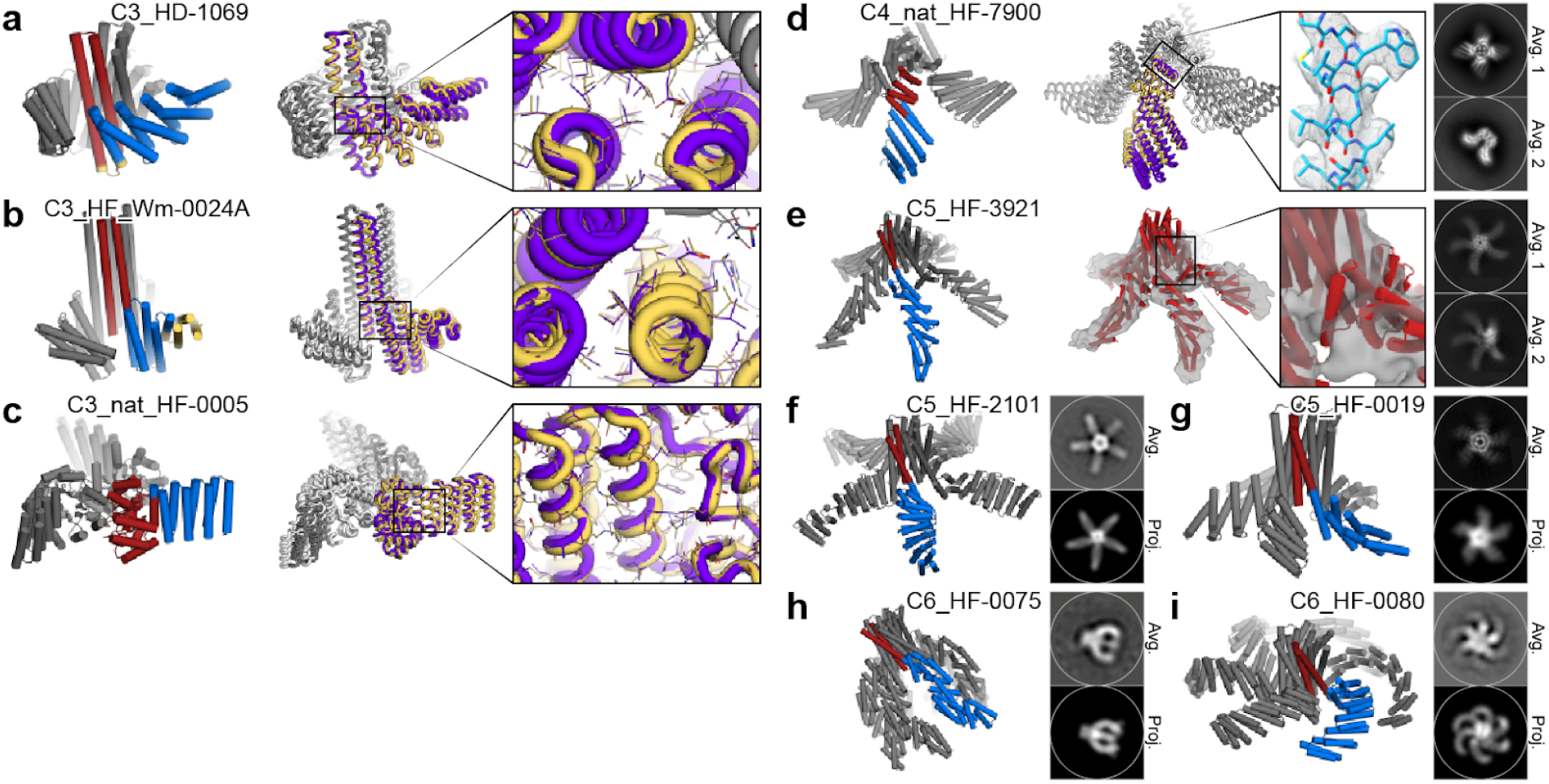
Homo-oligomer diversification by repeat protein fusion. Central oligomer units are shown in red and fused DHRs in blue. Design of (a) C3_HD-1069 (designed loop shown in yellow), (b) C3_HF_Wm-0024A (additional WORMS fusion shown in yellow), and (c) C3_nat_HF-0005. Overlay of the design model (purple/grey) and crystal structure (yellow/white) shows the overall match of the backbone. Inset shows the correct placement of the rotamers in the designed junction region. Design of higher order oligomer fusions (d) C4_nat_HF-7900 and (e) C5_HF-3921 as characterized by cryo-EM. C4_nat_HF-7900 design model (purple/grey) and Cryo-EM map (yellow/white), with inset highlighting the high resolution (~3.8Å) density. C5_HF-3921 inset showing density surrounding the designed junction. (f) C5_HF-2101, (g) C5_HF-0019, (h) C6_HF-0075, and (i) C6_HF-0080 showed good overall match to its negative-stain EM 2D class averages (top) from one direction; predicted projection map for comparison on the bottom.

#### HelixFuse (HF) approach

The same set of DHRs and HBs were combinatorially fused together by overlapping the terminal helix residues in both directions (“AB”: c-terminus of HB to n-terminus of DHR, “BA”: n-terminus of HB to c-terminus of DHR)^17^. On the HB end, up to 4 residues were allowed to be deleted to maximize the sampling space of the fusion while maintaining the structural integrity of the oligomeric interface. On the DHR end, deletions up to a single repeat were allowed. After the C-beta atoms are superimposed, a RMSD check across 9 residues was performed to ensure that the fusion results in a continuous helix. If no residues in the fused structure clash (Rosetta centroid energy < 10), sequence design was carried out at all positions within 8Å of the junction. This first step of the fusion sampling is wrapped into the Rosetta MergePDBMover^17^. After sequence design around the junction region^14,28^, fusions were then evaluated based on the number of helices interacting across the interface (at least 3), buried surface (sasa > 800) across the junction, and shape complementarity (sc > 0.6) to identify designs likely to be rigid across the junction point. In total, the building block library generated *in silico* by HelixFuse using HB hubs and DHR arms in this set consists of 490 C2s, 1255 C3s, 107 C5s, and 87 C6s.

As a proof of concept, select fusions to C5 (5H2LD-10^7^) and C6 (6H2LD-8) (in press) helical bundles were tested experimentally, as structures of higher cyclic symmetries were historically more difficult to design thus resulting in a lack of available scaffolds. Contrarily, larger structures are easier to experimentally characterize via electron microscopy due to their size. A total of 65 designs whose genes encoding the designs were synthesized and subsequently expressed in *E.coli*, 45 were soluble, and 23 were monodisperse by SEC. Of the ones that were selected for SAXS analysis, 7 had matching SAXS profiles (Figure S4-5). Cryo-electron microscopy of C5_HF-3921 followed by 3D reconstruction showed that the positions of the helical arms are close to the design model (Figure 2e, Figure S8 & 9). By negative-stain electron microscopy (EM), C5_HF-2101, C5_HF-0019, C6_HF-0075, and C6_HFuse-0080 (Figure 2f-i respectively) were class averaged and the top-down view clearly resembles that of the designed model and its predicted projection map (Figure S10, S12, S13, and S14 respectively). From negative-stain EM class averaging, off-target states can sometimes be observed; most obvious in C5_HF-0007 (Figure S11) and C6_HF-0075 (Figure S13), and less in C5_HF-0019 (Figure S12), where in some cases an incorrect number of DHR arms can be observed in the 2D class averages.

We also applied the method to two non-helical bundle oligomers – 1wa3, a native homo-trimer^29^ and tpr1C4_pm3, a designed homo-tetramer^25^. As described above, we fused DHRs to the N-terminal helix of 1wa3 and the C-terminal helix of tpr1C4_pm3. For 1wa3, from the 13 designs were expressed for experimental validation, 10 displayed soluble expression and showed clean monodispersed peaks by SEC. Through X-ray crystallography, we were able to solve C3_nat_HF-0005 to 3.32Å resolution (Figure 2c). A total of 16 tpr1C4_pm3 fusions were tested, 14 found to be soluble, and 10 displayed monodispersed peaks by SEC. The best behaving designs were analyzed by electron microscopy. C4_nat_HF-7900 was found to form monodisperse particles by cryo EM, with the 3D reconstruction modeled to 3.7Å resolution (Figure 2d, Figure S5-S7). Both the crystal structure of C3_nat_HF-0005 and the model of the cryo-EM reconstruction of C4_nat_HF-7900 show very good matches near the oligomeric hub of the protein where side chains are clearly resolved and as expected. However, it can be seen that they deviate from the design model at the most distal portions of the structure. This is likely due to the inherent flexibility of the unsupported terminal helices of the DHRs^17,23,30^ and lever arm effects which increase with increasing distance from the fusion site (Figure S15).

To extend the complexity of structures that can be generated, we built libraries of heteromeric two chain building block by fusing repeat proteins to two hetero-dimeric helical bundles (DHD-13, DHD-37)^21^ (Figure 1a). The fusion steps are identical, except for an additional step of merging the chain A and chain B fusions and checking for clashes and incompatible residues. In total, 2740 heterodimers were generated *in silico* to be part of the library. While the homo-oligomeric fusions are good building blocks for objects with higher order point group symmetries, hetero-oligomeric fusions are needed at segments without symmetry, such as building cyclic structures and/or connecting different axes of symmetry in higher order architectures (described below).

With a sufficiently high design success rate, the individual oligomers do not need to be experimentally verified before being used to build larger structures. Since all building blocks terminate in repeat proteins which can be fused anywhere along their length, the total number of possible three building block fusions which can be built from this set is extremely large, which could offset the degree of freedoms lost to symmetry constraints. The combined library for higher order oligomers consists of both HelixDock and HelixFuse generated building blocks; overall, the HelixFuse structures tended to have smaller interfaces across the junction, and thus less overall hydrophobicity than those generated by HelixDock. While the HelixFuse are less globular than their HelixDock counterparts, the smaller interface may contribute to the higher fraction of designs being soluble (~70% vs ~55%). The HelixDock method also requires an additional step of building a new loop between the HB and DHR, which is another potential source of modeling error, and takes significantly more computational hours. Overall, the final fraction with single dominant species in SEC traces (examples shown in Figure S1-S5) profiles are similar (~35%).

### Section II: Assembly of higher order symmetric structures from repeat protein-helical bundle fusion building blocks

To generate a wide range of novel protein assemblies without interface design, we took advantage of the protein interfaces in the library of building blocks described in the previous section, which are oligomers with repeat protein arms. Assemblies are formed by splicing together alpha helices of the repeat protein arms in different building blocks. In our implementation, the user specifies a desired architecture and the symmetries and connectivity of the constituent building blocks. The method then iterates through splices of all pairs of building blocks at all pairs of (user specified, see methods) helical positions; this very large set is filtered on the fly based on the rms of the spliced helices, a clash check, off-architecture angle tolerance, residue contact counts around splice, helix contact count, and redundancy; all of which can be user specified parameters (see methods). The rigid body transform associated with each splice passing the above criteria is computed; for typical pairs of building blocks allowing 100 residues, 100 x 100 = 10,000 unfiltered splices are possible.

Assemblies of these building blocks are modeled as chains of rigid bodies, using the transform between coordinate frames of entry and exit splices, as well as transform between entry splice and coordinate frames of the building blocks. Assemblies are built, in enumerative fashion or with monte carlo, by simple matrix multiplication. For efficiency, only prefiltered splices are used. This technique allows billions of potential assemblies to be generated per cpu hour. Criteria for a given assembly design problem can include any operation defined on the rigid body positions of the building blocks. In this work, we use the transform from the start and end building blocks. To form Cx cyclic oligomers, the rotation angle of the transform must be 360/x, and the translation along the rotation axis must be zero. To form tetrahedral, octahedral, icosahedral, and dihedral point group symmetries from cyclic building blocks, the symmetry axes of the start and end building blocks must intersect, and form the appropriate angle for the desired point group; for example, a 90° angle creates dihedral symmetry.

This rapid symmetric architecture assembler through building block fusion has been implemented in a program called WORMS (Wm) which provides users with considerable control over building block sets, geometric tolerances, and other parameters and enables rapid generation of a wide range of macromolecular assemblies. The desired architecture is entered as a config file (or command line option) in the following format illustrated for a 3-part fusion with icosahedral symmetry:

~~~
[‘C3_N’, orient(None, ‘N’)), (‘Het:CN’, orient(‘C’, ‘N’)), (‘C2_C’, orient(‘C’, None)]
Icosahedral(c3=0, c2=−1)
~~~

The architecture is specified first, here an icosahedral structure constructed from a C3 and a C2 building block, and then how the selected building blocks types from the loaded databases are to be linked together (like a worm). In this example, a C3 building block with an available N-terminus ‘C3_N’ is to be fused to a hetero-dimeric building block ‘Het:CN’ via an available C-terminus, and the N-terminus of the same ‘Het:CN’ is in turn to be fused to a third C2 building block ‘C2_C’ through an available C-terminus. The ‘None’ designation marks that there are no additional unique connections to be made on that segment. Through the assignment of ‘c3=0’ and ‘c2=−1’, the first and last building blocks are declared as the C3 component and the C2 component, respectively.

The building blocks are cached the first time they are read in from the database files, which can range from a single entry per type to thousands, due to the combinatorial nature of the first fusion step. See supplementary information for more details regarding additional options, architecture definitions, and database syntax. With hundreds to thousands of building blocks each with ~100 residues available for fusion, the total number of three way fusions is on the order of greater than 10^14^, so optimization of efficiency in both memory usage and CPU requirements was critical in WORMS software development.

Once building block combinations are identified that generate the designed architecture (within a user specifiable tolerance), explicit atomic coordinates are calculated and used for clash checking, redundancy filtering, and any other filtering that requires atomic coordinates. Models for each assembly passing user specified tolerances are constructed in Rosetta, scored and output for subsequent sequence design.

#### Generation of cyclic “crowns” (Crn)

We generated C3, C4, and C5 assemblies with WORMS using two designed heterodimer fusions from HelixFuse, as described above. This resulted in head-to-tail cyclic ring structures (Figure 3a), generated by the following configuration (C3 as an example):

~~~
[(‘Het:CN’, orient(None, ‘N’)),(‘Het:CN’, orient(‘C’, None))]
Cyclic(3)
~~~

**Figure 3.**
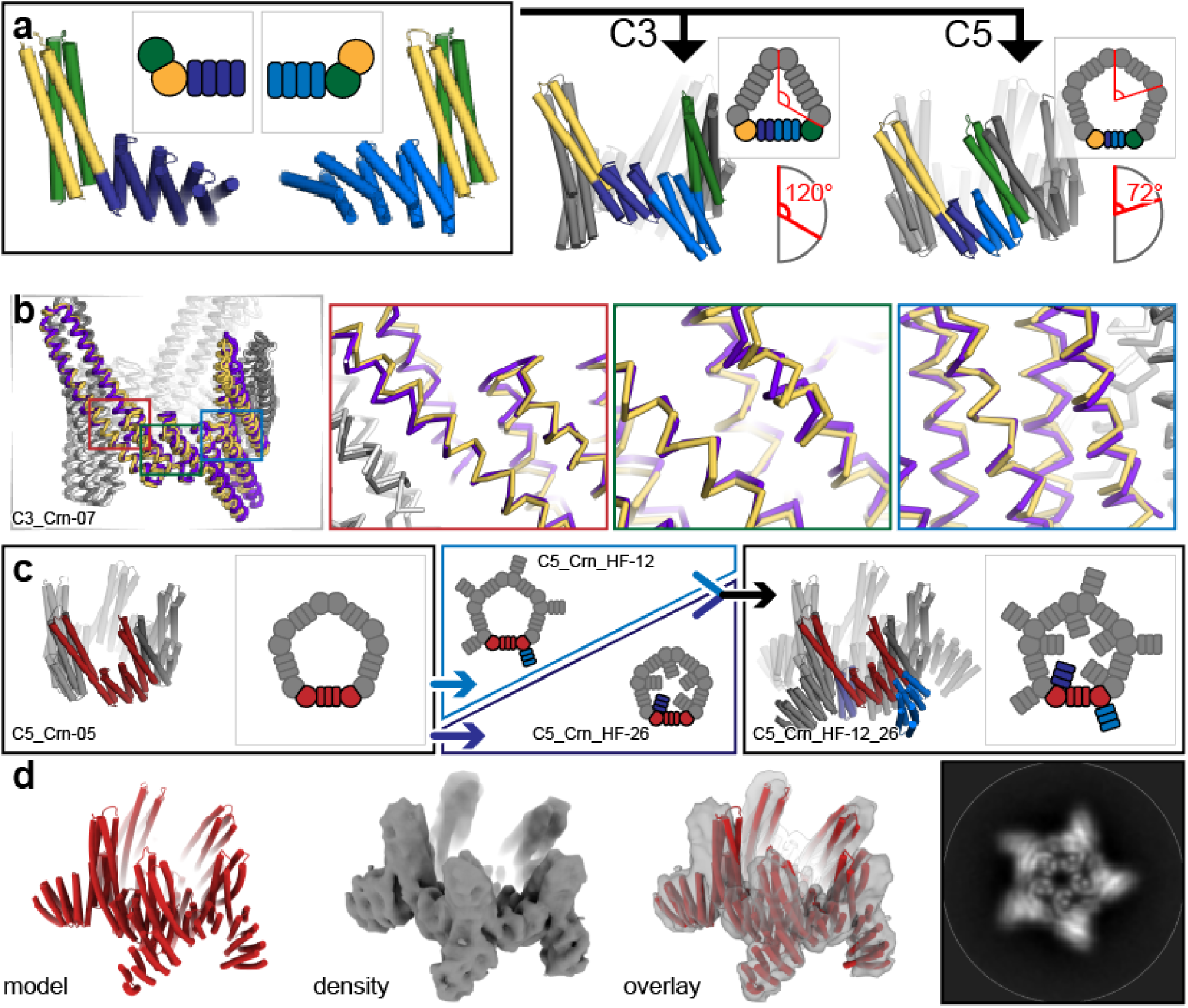
Design of cyclic “crown” (Crn) structures from heterodimeric building blocks. (a) Hetero-dimeric HB (green/yellow) fused with different DHRs (shades of blue) were fused together using WORMS by enforcing a specific overall cyclic symmetry (C3 and C5 shown). (b) The backbones of the crystal structure (yellow/white) of C3_Crn-05 overlaid with the design model (purple/grey). Insets show the backbone matching focused at each of the fusion locations. (c) A C5 crown (C5_Crn-07, asymmetric unit in red) was fused to DHR units on either exterminal (“C5_Crn_HF-12”, blue arrow) or internal termini (“C5_Crn_HF-26”, dark blue arrow). The two structures were then merged together to generate a double fusion (“C5_Crn_HF-12_26”, black arrow). (d) Cryo-EM class average of the fused 12_26 structure; the major C5 species shown. 3D reconstruction shows the main features of the designed structure are present, as is also evident in the class average (right).

Following fusion, the junction residues were redesigned to favor the fusion geometry and filtered as above. Seven C3s, seven C4s, and eight C5s were selected and tested experimentally. All yielded soluble protein, and 6, 2, and 1 respectively showed a single peak at the expected elution volume via SEC. We solved the structure of the C3_Crn-05 to 3.19A resolution (Figure 3b). The overall topology is as designed and the backbone geometry at each of the three junctions is close to the design model. A deviation at the tip of the undesigned heterodimeric HB is likely to due to crystal packing. C5_Crn-07 chromatographed as a single peak by SEC and was found to be predominantly C5 by negative-stain EM (Figure 3d), but minor off-target species (C4, C6, and C7) were also observed (Figure S16). Each of these structures experimentally verifies three distinct helical fusions (two HelixFuse, one WORMS) from a previously unverified building block library.

To further increase the diversity of the crown structures, we recursively ran HelixFuse on both termini of C5_Crn-07 (Figure 3c). Six (6) N-terminal and 24 C-terminal fusions were selected and experimentally tested. All were soluble, but had large soluble aggregate fractions when analyzed by SEC. When the peaks around the expected elution volumes were analyzed by negative-stain EM, ring-like structures were found in many of the samples. To facilitate EM structure determination, we combined a c-terminal fusion (C5_Crn_HF-12) and an n-terminal (C5_Crn_HF-26) fusion to generate C5_Crn_HF-12_26 (Figure 3c), which resulted in a much cleaner and monodisperse SEC profile (Figure S17). Cryo-electron microscopy of 12_26 revealed the major population of C5 (77%) structures in addition to C4 (1%), D5 (8%), and C6 (12%) subpopulations (Figure S17). We hypothesize that the D5 structure is due to transient interactions of histidines placed on the loops for protein purification. The final 3D reconstruction to 5.6Å resolution shows that the major characteristics of the design model are present, despite some splaying of the undesigned portion of the heterodimeric HB relative to the design model (Figure 3d).

#### Generation of two-component dihedral assemblies

Dihedral symmetry protein complexes are attractive building blocks for making higher order 2D arrays and 3D crystal protein assemblies, and can be useful for receptor clustering in cellular engineering^31^. We first set out to design dihedral protein assemblies of D2 symmetry. A set of C2 homo-oligomers with DHR termini (described above) were fused with select *de novo* hetero-dimers (tj18_asym13, unpublished work) using WORMS (schematics shown in Figures 4a-b). The D2 rings harbored total 8 protein chains with 2 chains (two-component) as the asymmetric unit. To generate these rings, we used a database of building blocks containing 7 homo-dimers and 1 heterodimer using the following configuration:

~~~
[(‘C2_C’, orient(None, ‘C’)), (‘Het:CN’, orient(‘N’, ‘C’)), (‘C2_N’, orient(‘N’, None))]
D2(c2=0, c2b=−1)
~~~

**Figure 4.**
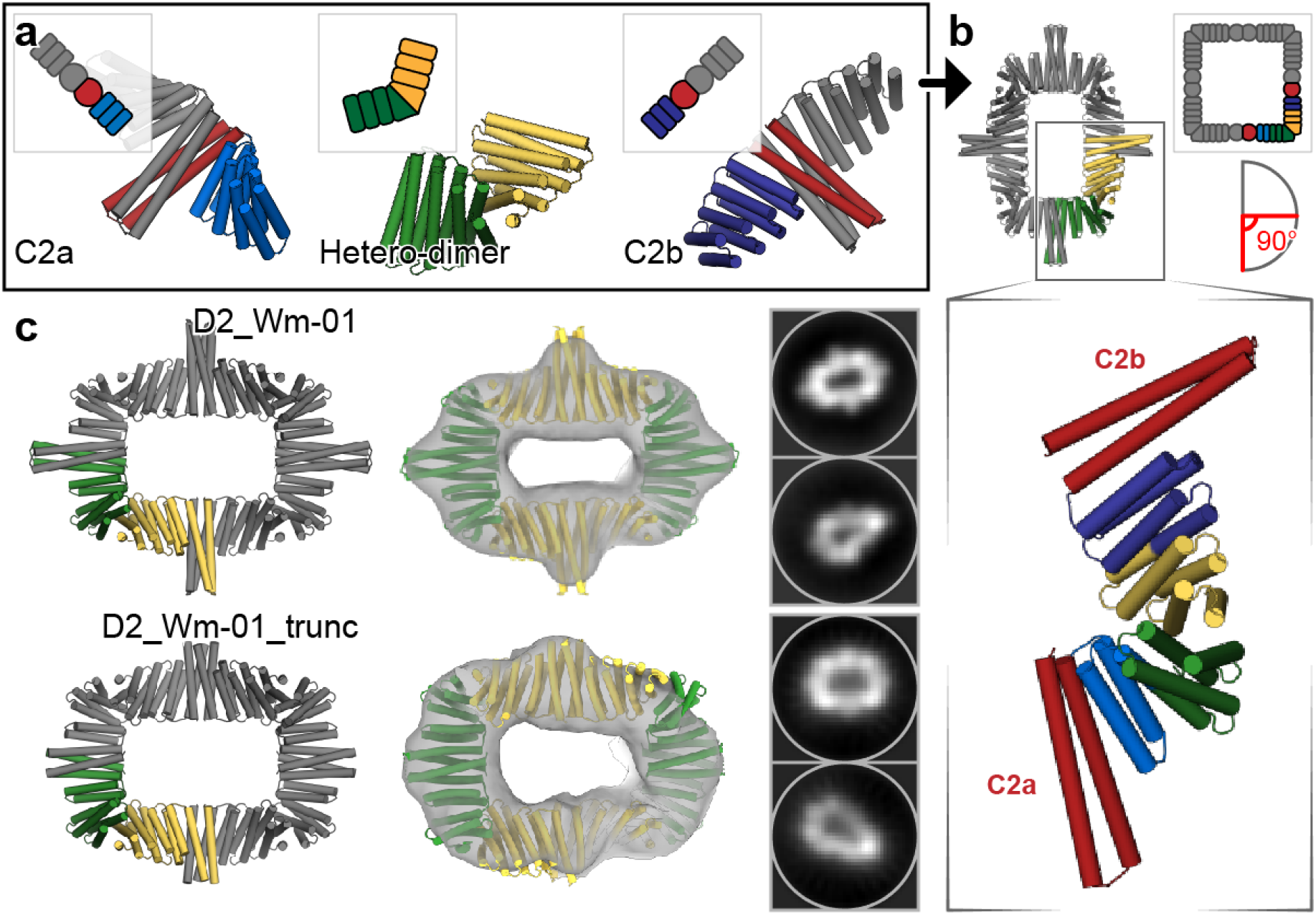
Design of two-component dihedral rings using WORMS (Wm). (a) Two different homodimeric HBs (red) with DHR extensions (shades of blue) were aligned to their respective symmetrical axes with dihedral symmetry. An additional heterodimer (green/yellow) was placed between them and systematically scanned and fused together to design an 8-chain D2 ring. (b) The final asymmetric unit shown in green/yellow while the inset preserves the original colors. (c) Negative-stain EM followed by 2D average and 3D reconstruction of D2_Wm-01 and D2_Wm-01_trunc show that the major features of the designs were recapitulated (left) designed model, (middle) overlay of the designed models with the 3D reconstructions, (right) 2D averages.

Of 208 outputs, we selected 6 designs to test, out of which three expressed as soluble two-component protein assemblies as indicated by Ni-NTA pulldown and subsequent SDS-PAGE experiments. Of these, two designs (designated as D2_Wm-01 and D2_Wm-02) eluted as expected by SEC and had SAXS profiles that matched with the designed models (Figure S18 & 19).

To characterize the structures of D2_Wm-01 and D2_Wm-02 in more detail, we performed negative-stain EM and subsequent 2D averaging and 3D refinement. 2D averaging shows the resemblance of the designed model with the experiment-determined structures, whereas 3D refinement indicated accurate design of D2_Wm-01 and D2_Wm-02 at ~16 Å resolution (Figure 4c, Figure S19).

The homo dimeric building blocks used in D2_Wm-01 and D2_Wm-02 have large interface areas (~35 residues long; 5 heptads). We sought to reduce the interface area by truncating the helices to facilitate expression of the components and reduce off target interactions. Deletion of one heptad from either of the homodimers of D2_Wm-01 (designated D2_Wm-01_trunc) resulted in a single and much narrower SEC peak of the expected molecular weight (Figure S18). Negative-stain EM followed by 2D averaging and 3D refinement indicated monodispersed particles with accurate structure as of the designed model (Figure 4c).

#### Generation of one-component tetrahedral protein cages

Idealized ankyrin homo-dimers^25^ based on ANK1 and ANK3 and selected HBs^20^ were combined to design one-component tetrahedral cages capable of hosting engineered DARPIN binding sites. For each combination, a monomeric ankyrin that perfectly matches the homo-dimer backbone was added as a spacer in between the homo-oligomers, thus extending the ankyrin homo-dimer by several repeats (Figure 5a). To set up this architecture, the following configuration can be used:

~~~
[(‘C2_N’, orient(None, ‘N’)), (‘Monomer’, orient(‘C’, ‘N’)), (‘C3_C’, orient(‘C’, None))]
Tetrahedral(c2=0, c3=−1)
~~~

**Figure 5.**
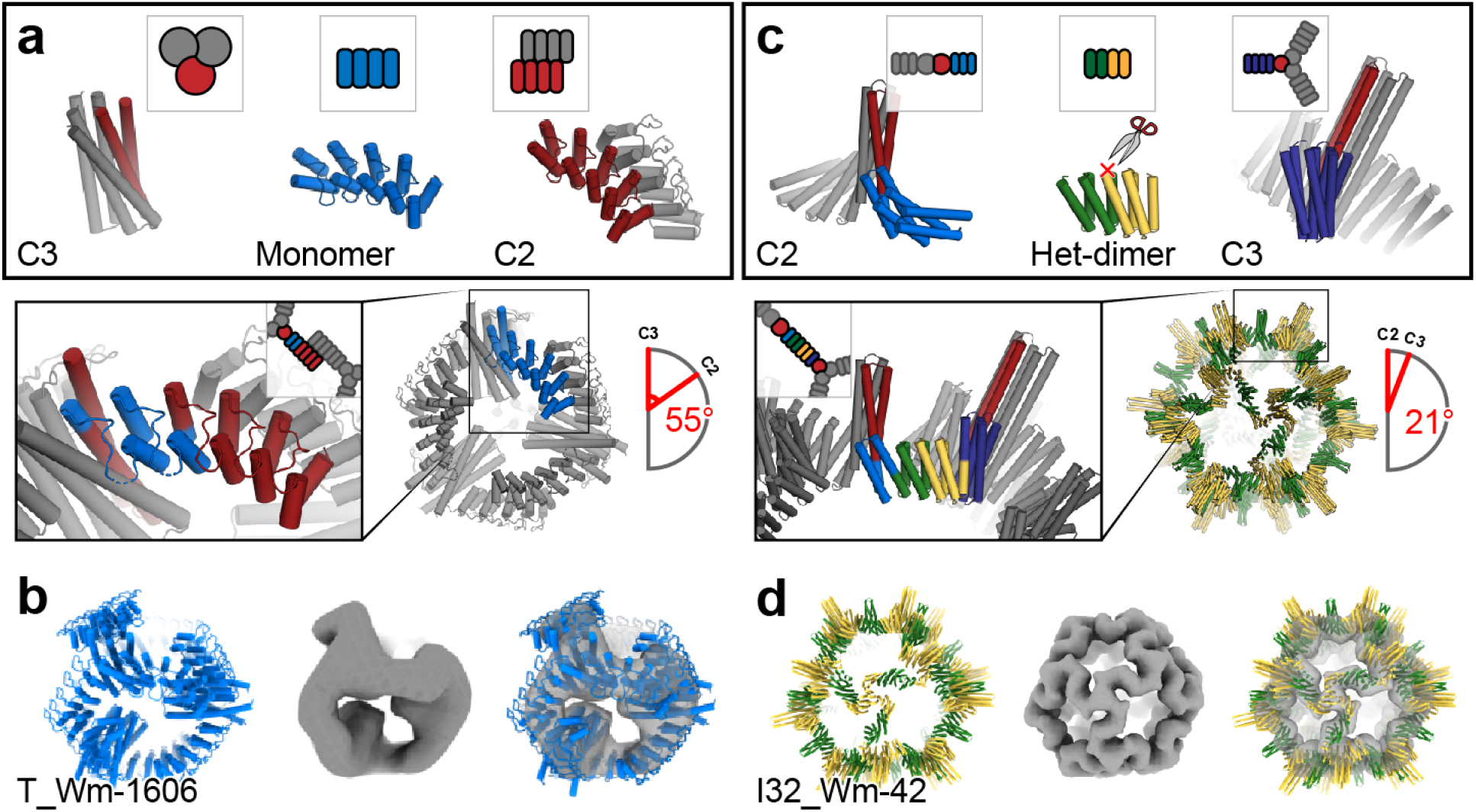
Design of assemblies with point group symmetry through helical fusion with WORMS. (a) Tetrahedron design schematic. A HB and a C2 homo-oligomeric made from ankyrin repeat proteins were aligned to their respective tetrahedral symmetry axis (red), and connected via fusion to Ankyrin repeat monomers (blue) to generate the target architecture. (b) 3D reconstruction reveals a well fitting map of T_Wm-1606. (c) Icosahedral design schematic. Libraries of unverified cyclic fusion homo-dimers and trimers were aligned to the corresponding icosahedral symmetry axes. Using WORMS, fusions to DHRs split in the center that hold the two homo-oligomers in the orientations which generate icosahedral structures were identified. (d) Cryo-EM 3D reconstruction of I32_Wm-42 closely matches the designed model.

Due to the relatively small space of possibilities because of the limited building block set, only 27 valid fusion combinations were identified, of which 20 involved ankyrin homo-dimer extension at its N-terminus and the remaining 7 at its C-terminus. Eight (8) were selected by manual inspection for further sequence design at fusion regions and experimental characterization.

All 8 constructs were expressed and two were found to be soluble with mono-disperse elution profile peaks by SEC. The two promising structures were very similar, containing different helical bundles whose backbone geometry was identical, but with different internal hydrogen-bond networks. As the two were so similar, only one (T_Wm-1606) was selected for negative-stain EM and discrete particles were observed whose 2D class averages and 3D reconstruction to 20Å matched the computational model (Figure 5b). There was also good agreement between experimental SAXS profiles and profiles computed from the design model (Figure S20).

#### Generation of two-component icosahedral protein cages

Point group symmetry nanocages have been successfully designed using docking followed by interface design^5–7^. To build such structure using our building blocks with the smaller and weaker interfaces that give rise to cooperative assembly^32–34^, we systematically split each DHR at the loop in the center of four repeats, resulting in a hetero-dimeric structure with two repeats on each side. The resulting interfaces are considerably smaller than in for example our *de novo* designed helical bundles. The WORMS protocol was then applied using the C5, C3, and C2 HelixFuse libraries described above at their corresponding tetrahedral, octahedral, and icosahedral symmetry axes. The split DHRs were then sampled to be connected in the center to each of the two symmetrical oligomers (Figure 5c), using the configuration described above. Following fusion, sequence design was performed at each of the two new junctions.

57 total designs were selected for experimental characterization; 25 co-eluted by Ni-NTA chromatography, and of these 7 designs had large peaks in the void volume in SEC chromatography as expected for particles of this size. When the peaks were collected and re-analyzed with a Sephacryl 500 column, one design, I32_Wm-42 (icosahedral architecture) was resolved into a void and a resolved peak (Figure S21). Cryo-EM analysis of the resolved peak reveals well formed particles that when reconstructed to 9Å resolution, accurately match the design model, including the distinct “S” shaped turn between the C3 and C2 axes (Figure 5d). This structure is considerably more open than previous icosahedral cages built by designing non-covalent interfaces between homo-oligomers. For another design, T32_Wm-24, while cage was not formed, we were able to crystallize the polar-capped trimer component (C3_HF_Wm-0024A) and solve the structure by x-ray diffraction to 2.69Å (Figure 2B). The structure clearly shows that both of the newly designed junctions (from HelixFuse and WORMS) are as designed, matching the design model.

The 120 subunit I32_Wm-42 icosahedral nanocage has a molecular weight of 3.4 MDa and a diameter of 42.7 nm and illustrates the power of our combined hierarchical approach. I32_Wm-42 is constructed from five building blocks (two helical bundles and three repeat proteins) combined via four unique rigid junctions; the EM structure demonstrates that all were modeled with reasonable accuracy. The combination of the HelixDock and HelixFuse helix fusion methods created a large set of over 1500 oligomeric building blocks from which WORMS was able to identify combinations and fusion points that generated the icosahedral architecture; this example is notable because none of the oligomeric building blocks had been previously characterized experimentally. With fewer unknowns, either using less segments or a larger fraction of previously validated building blocks, we expect considerable improvement of the overall success rate.

## Discussion

Our general rigid helix-fusion based pipeline fulfills the promise of early proposals^16,35^ in providing a robust and accurate procedure for generating large protein assemblies by fusing symmetric building blocks and avoiding interface design, and should streamline assembly design for applications in vaccine development, drug delivery and biomaterials more generally. The set of structures generated here goes considerably beyond our previous work with rigid helical fusions^18^, and the “WORMS” software introduced here is quite general and readily configurable to different nanomaterial design challenges. WORMS can be easily extended to other symmetric assemblies including 2D arrays and 3D crystals, and should be broadly useful for generating a wide range of protein assemblies.

DNA nanotechnology has had advantages in modularity and simplicity over protein design because the basic interactions (Watson-Crick base pairing) and local structures (the double helix) are always the same. Proteins in nature exhibit vast diversity compared to duplex DNA, and correspondingly, re-engineering naturally occuring proteins and designing new ones has been a more complex task than designing new DNA structures. The large libraries of “clickable” building blocks--helical bundle – repeat protein fusions-- and the generalized WORMS software for assembling these into a wide range of user specifiable architectures that we present in this paper are a step towards achieving the modularity and simplicity of DNA nanotechnology with protein building blocks. Although this modularity comes at some cost in that the building blocks are less diverse than proteins in general, they can be readily functionalized by fusion to protein domains with a wide range of functions. We show that it is possible to genetically fuse DHR “adapters” to natural proteins; these proteins can then be used in larger assemblies through WORMS with less likelihood of disrupting the original protein fold. Proteins of biological and medical relevance (binders like protein A, enzymes, etc.) can be used as components and combined with *de novo* designed HBs and DHRs to form nanocages and other architectures.

Moving forward, there are still a variety of challenges to address. The larger the set of building blocks for WORMS the more precisely the geometric constraints associated with the desired architecture can be achieved, and hence it is advantageous to use the very large *in silico* libraries of building blocks that can be created by helical bundle – repeat protein fusion rather than the very much smaller sets of fusions that can be experimentally characterized in advance (tens of thousands compared to tens). It will be important to understand how uncertainties in the structures of the *in silico* fusions translate into uncertainties in the structures of the resulting architectures, and more generally, how to further improve the fusion approach so that the *in silico* structures are nearly perfectly realized. As the assemblies become more complex with different building blocks and total number of subunits, more alternative structures become possible. Understanding how to achieve cooperative assembly and controlling for specificity of the desired assembly over alternatives will be an increasingly important challenge as the complexity of the target nanomaterials increases.

### Computational Methods Summary

#### RosettaRemodel Forward Folding

To test the extent to which the designed sequences encode the designed structure around the junction site, we used large scale *de novo* folding calculations. Due to computational limitations with standard full chain forward folding^36,37^, we developed a similar but alternate approach for larger symmetric structures. Using RosettaRemodel^27^ in symmetry mode (reversing the anchor residue for cases where the helical bundle was at the C-terminus), we locked all residues outside the junction region as rigid bodies, only allowing 40 residues starting from the end of the HB in the primary sequence direction of the DHR to be re-sampled. The blueprint file was set up to be agnostic of secondary structure in this segment of protein and we deleted all DHR residues past the first two helices after the rigid body region to reduce CPU cost. Each structure was set to at least 2000 trajectories to create a forward folding funnel.

#### WORMS

The WORMS software overall requires two inputs, a database of building block entries (format described in Supplementary Information in detail) and a configuration file (or command line options) as described in the main text to govern the overall architecture. While some segments can be of single building blocks of interest, to generate a wide variety of outputs, tens to thousands of entries per segment should be used. The number of designs generated also depends on the number of fusion points allowed, as the size of the space being sampled increases multiplicatively with the number of segments being fused. There are many options available to the user to control the fusions which are output as solutions; we have tuned the default options to be relatively general-use (see Supplementary Information for description of options). A key parameter is the *tolerance*, he allowed deviation of the final segment in the final structure away from its target position given the architecture. For different geometries the optimal values vary; for example the same tolerance values involve more drastic error in icosahedral symmetry than cyclic symmetry. The WORMS code is specifically designed to generate fusions that have a protein core around the fusion joint; unless specified using the *ncontact_cut, ncontact_no_helix_cut*, and *nhelix_contacted_cut* option set, the code will not produce single extended helix fusions.

### Brief Experimental Methods

#### Gene preparation

All amino acid sequences derived from Rosetta were reverse translated to DNA sequences and placed in the pET29b+ vector. For two-component designs, all designs were initially constructed for bi-cistronic expression by appending an additional ribosome binding site (RBS) in front of the second sequence with only one of the components containing a 6xHis tag. Genes were synthesized by commercial companies: Integrated DNA Technologies (IDT), GenScript, Twist Bioscience, or Gen9.

#### Protein expression and purification

All genes were cloned into *E. coli* cells (BL21 Lemo21 (DE3)) for expression, using auto-induction^38^ at 18° or 37°C for 16-24 hours in 500mL scale. Post-induction, cultures were centrifuged at 8,000xG for 15 minutes. Cell pellets were then resuspended in 25-30mL lysis buffer (TBS, 25mM Tris, 300mM NaCl, pH8.0, 30mM imidazole, 0.25mg/mL DNase I) and sonicated for 2 minutes total on time at 100% power (10 sec on/off) (QSonica). Lysate was then centrifuged at 14,000xG for 30 minutes. Clarified lysates were filtered with a 0.7um syringe filter and put over 1-4mL of Ni-NTA resin (QIAgen), washed with wash buffer (TBS, 25mM Tris, 300mM NaCl, pH8.0, 60mM imidazole), then eluted with elution buffer (TBS, 25mM Tris, 300mM NaCl, pH8.0, 300mM imidazole). Eluate was then concentrated with a 10,000 m/w cutoff spin concentrator (Millipore) to approximately 0.5mL based on yield for SEC.

D2 proteins went through an extra round of bulk purification. Concentrated protein was heated at 90 °C for 30 minutes to further separate bacterial contaminants. Samples were then allowed to cool down to room temperature and any denatured contaminants were removed by centrifuging at 20,000xG.

#### Size exclusion chromatography (SEC)

All small oligomers were passed through a Superdex200 Increase 10/300 GL column (Cytiva) while larger assemblies were passed through a Superose 6 Increase 10/300 GL column (Cytiva) on a AKTA PURE FPLC system. The mobile phase was TBS (TBS, 25mM Tris, 300mM NaCl). Additionally, for the icosahedral assembly, an additional custom packed 10/300 Sephacryl500 column (Cytiva) was used to separate out the void. Samples were run at a speed of 0.75mL/min and eluted with 0.5mL fractions.

#### Protein Characterization

See supplementary information for detailed methods regarding SAXS sample preparation, electron microscopy, and x-ray crystallography.

## ACKNOWLEDGMENTS

This work was supported by the National Science Foundation (NSF) award 1629214 (DB), a generous gift from the Audacious Project, the Open Philanthropy Project Improving Protein Design Fund, the National Institute of General Medical Sciences (R01GM120553 to D.V.), the National Institute of Allergy and Infectious Diseases (HHSN272201700059C to DV), a Pew Biomedical Scholars Award (DV), an Investigators in the Pathogenesis of Infectious Disease Award from the Burroughs Wellcome Fund (DV) and the University of Washington Arnold and Mabel Beckman cryo-EM center. YH was supported in part by a NIH Molecular Biology Training Grant (T32GM008268). RM is a recipient of the Washington Research Foundation (WRF) Innovation fellowship and his research is funded in part by the US DOE BES Energy Frontier Research Center CSSAS (The Center for the Science of Synthesis Across Scales) located at the University of Washington (award number DESC0019288). UN was supported in part by PHS NRSA (T32GM007270) from NIGMS. AC is a recipient of the Human Frontiers Science Program Long Term Fellowship and a Washington Research Foundation Senior Fellow.

This work was conducted at the Advanced Light Source (ALS), a national user facility operated by Lawrence Berkeley National Laboratory on behalf of the Department of Energy, Office of Basic Energy Sciences, through the Integrated Diffraction Analysis Technologies (IDAT) program, supported by DOE Office of Biological and Environmental Research. Additional support comes from the National Institute of Health project ALS-ENABLE (P30 GM124169) and a High-End Instrumentation Grant S10OD018483.

We thank staff at Advanced Photon Source beamline NE-CAT 24-ID-E for data collection. Northeastern Collaborative Access Team beamline supported by NIH grants P30GM124165 and S10OD021527, and DOE contract DE-AC02-06CH11357. We also want to thank Banumathi Sankaran at the Advanced Light Source (ALS) beamline 8.2.2 at Lawrence Berkeley National Laboratory for data collection. The Berkeley Center for Structural Biology is supported in part by the National Institutes of Health (NIH), National Institute of General Medical Sciences, and the Howard Hughes Medical Institute. The Advanced Light Source (ALS) is supported by the Director, Office of Science, Office of Basic Energy Sciences and US Department of Energy under contract number DE-AC02-05CH11231.

We thank Kristen Dancel-Manning and Alice Liang of the NYU Microscopy Laboratory, William Rice and Bing Wang of the NYU Cryo-EM Laboratory, Kashyap Maruthi, Ed Eng, Laura Yen, and Misha Kopylov of the New York Structural Biology Center, and members of the Bhabha/Ekiert labs for assistance with grid screening and data collection and helpful discussions. We especially thank Nicolas Coudray of the Bhabha/Ekiert lab for helpful discussions and guidance regarding EM data processing. Some of this work was performed at the Simons Electron Microscopy Center and National Resource for Automated Molecular Microscopy located at the New York Structural Biology Center, supported by grants from the Simons Foundation (SF349247), NYSTAR, and the NIH National Institute of General Medical Sciences (GM103310) with additional support from Agouron Institute (F00316), NIH (OD019994), and NIH (RR029300).

We would like to thank the Rosetta@Home user base for donating their computational hours to run our forward folding simulations. Thanks to George Ueda for the unpublished tj18_asym13 heterodimer. An additional thanks to Robby Divine and Josh Lubner for support in the documentation and development WORMS.

## AUTHOR CONTRIBUTIONS

YH, RM, and DB wrote the manuscript. YH, WS, and TB developed the HelixDock protocol; YH, RM, NIE, IV, and UN made designs and characterized experimentally with assistance from ET, AS, and CMC in protein production. YH and TB developed the HelixFuse protocol. IV developed the helical fusion method in .NET; WS and DB implemented it into the WORMS protocol; YH and RM assisted developing in its application. UN and ET crystallized and MJB solved the C3_HD-1069 structure. AK crystallized C3_nat_HF-0005, C3_HF_Wm-0024A, and C3_Crn-05. AB solved the crystal structure for C3_nat_HF-0005 and C3_HF_Wm-0024A. MJB solved the structure for C3_Crn-05. RLR, assisted by DE and GB, performed negative-stain and cryo-EM for all HelixFuse structures presented. YH designed and characterized crown structures and icosahedral cage; RM the dihedral structures, IV the tetrahedral cage. RM and AC performed initial EM screening of dihedral, cyclic and icosahedral WORMS structures. YJP, assisted by DV, performed negative-stain and cryo-EM for all WORMS structures presented. DB guided the project.

## ONLINE CONTENT

Crystallography data:

C3_HD-1069 (**6XT4**)
C3_HF_Wm-0024A (**6XI6**)
C3_nat_HF-0005 (**6XH5**)
C3_Crn-05 (**6XNS**)

Electron microscopy data:

C4_nat_HF-7900 (**6XSS, EMD-22305**)
C5_HF_3921 (**EMD-22306**)
C5_Crn_HF-12_26 (**EMD-XXXX**)
D2_Wm-01 (**EMD-XXXX**)
D2_Wm-01_trunc (**EMD-XXXX**)
D2_Wm-02 (**EMD-XXXX**)
T_Wm-1606 (**EMD-XXXX**)
I32_Wm-42 (**EMD-XXXX**)

## ADDITIONAL INFORMATION

Supplementary files:

Supplementary information *.pdf
HelixDock sequence design *.xml, *.symdef
HelixDock loop closure *.xml
HelixFuse *.xml
WORMS sequence design *.xml, *.xml
Design models *.pdb (as zip files)

