## supplemental information for "Hierarchical design of multi-scale protein complexes by combinatorial assembly of oligomeric helical bundle and repeat protein building blocks"

[1]Department of Biochemistry, University of Washington, Seattle, WA 98195
[2]Institute for Protein Design, University of Washington, Seattle, WA 98195
[3]Biological Physics, Structure and Design Graduate Program, University of Washington, Seattle, WA 98195
[4]Molecular Engineering Graduate Program, University of Washington, Seattle, WA 98195
[5]Molecular and Cellular Biology Graduate Program, University of Washington, Seattle, WA 98195
[6]Medical Scientist Training Program, University of Washington, Seattle, WA 98195
[7]Howard Hughes Medical Institute, University of Washington, Seattle, WA 98195
[8]Department of Cell Biology and Skirball Institute of Biomolecular Medicine, New York University School of Medicine, New York, NY 10016
[9]Department of Microbiology, New York University School of Medicine, New York, NY 10016
[#]These authors have contributed equally

**THIS DOCUMENT CONTAINS:**

1) Supplementary methods
2) Supplementary figures
3) Supplementary tables
4) Supplementary protein sequences

### 1) SUPPLEMENTARY METHODS

#### HelixDock Details

HelixDock designs were first also attempted as “two-body hydrophobic” (2BH) designs, where the extra loop was not added to connect the helical bundle and repeat protein chain termini after a hydrophobic interface was designed between them. A couple dozen designs were tested in this fashion, but they all poor solubility or showed highly heterogeneous assemblies by SEC. This was hypothesized to be caused by the weak association of the repeat protein to the helical bundle; the final assembly was in constant equilibration and did not maintain the full stoichiometry. All further attempts with HelixDock were designed as “one-body hydrophobic” (1BH), where the helical bundle and repeat protein were closed into a single chain, as described in the main text. The HelixDock protocol is broken down into three distinct parts: docking, interface design, and loop closure.

#### HelixDock: Docking

The docking was performed using a modified version of the sicdock app, as previously described to generate cyclic oligomers from monomers. A new type of symmetry was added on to allow the docking of monomers in a symmetric manner (in this case, a repeat protein monomer) in all 6 degrees of freedom (dof) to a symmetry matching oligomer in the center that is not perturbed (in this case, a helical bundle). This final resulting architecture can be described as two matching cyclic symmetries stacked on top of each other along the z-axis; this definition will be used again as a symdef file during RosettaDesign. The docks were filtered based on their motif score, which is an estimate of the interface size and the likelihood of the dock to generate a decent interface post-design. An additional filter was used to make sure the termini between the helical bundle and repeat protein were compatible; the N- and C- terminus needed to be within 9 angstroms of one another.

**HelixDock: Sequence Design and Loop Closure**

To model the dock output correctly in Rosetta, we generated new symdef files which allowed the architecture as described above. Using SymDofMover, we were able to regenerate the docked conformation and select the relevant residues for interface design and subsequent filtering. To close the termini between the helical bundle and repeat protein, the ConnectChainsMover was used. After loop closure, residues at and around the new loop were redesigned and scored to ensure compatibility. Example *.sym and *.xml files that accomplishes these steps are available in the supplemental materials.

### WORMS

Full source code and repository can be found at:
https://github.com/willsheffler/worms

### WORMS relevant command line options:

The full list of available command line options can be initiated with --help.

#### I/O options

| Option | Default | Description |
|---|---|---|
| **--geometry** | | Specifies geometry (see architecture below). |
| **--bbconn** | | Specifies connectivity (see architecture below). |
| **--config_file** | | Specifies the config file to be used (see architecture below). |
| **--nbblocks** | 64 | The maximum number of building blocks that can be used in each segment. |
| **--dbfiles** | | Space delimited list of database files to be read in (see databases below). |
| **--shuffle_bblocks** | 1 | Uses a random set of building blocks instead of sequential from the top; relevant only if nbblocks < total actual bblocks available in that segment. |
| **--max_output** | 1000 | Maximum number of pdbs to output. |

#### Splice Level Filtering options

| Option | Default | Description |
|---|---|---|
| **--no_duplicate_bases** | 1 | Prevents duplicated 'bases' in the final structure (see databases below). |
| **--min_seg_len** | 15 | Minimum length required for each segment in the fusion. |
| **--splice_rms_range** | 5 | During splicing, this specifies the number of residues to check for rms at the fusion junction. (+/-) this many residues. |
| **--splice_max_rms** | 0.7 | Maximum rms allowed at the fusion junction. |
| **--splice_ncontact_cut** | 38 | Minimum number of contacts required across the interface. |
| **--splice_ncontact_no_helix_cut** | 6 | Minimum number of contacts required across the interface after removing the fusion helix. This filters against fusions where there are no additional interactions between the segments. |
| **--splice_nhelix_contacted_cut** | 3 | Minimum number of helices in contact after removing the fusion helix. |
| **--splice_max_chain_length** | 450 | Maximum final chain length after fusion. |
| **--tolerance** | 1.0 | Angstrom deviation from final structure, respective to its ideal axis. |

#### PyRosetta Level Filtering Options

| | | |
|---|---|---|
| **--max_score0** | 4.0 | Asymmetric Score0 filter after Rosetta scoring. |
| **--full_score0sym** | 1.0 | Symmetric Score0 filter after Rosetta scoring. |
| **--max_com_redundancy** | 4.0 | Computes the center of mass for each segment and filters designs out if the same building block is used at the same segments and their center of mass are in similar positions. |
| **--postfilt_splice_rms_length** | 9 | PyRosetta version of **--splice_rms_range** |
| **--postfilt_splice_max_rms** | 0.7 | PyRosetta version of **--splice_max_rms** |
| **--postfilt_splice_ncontact_cut** | 40 | PyRosetta version of **--splice_ncontact_cut** |
| **--postfilt_splice_ncontact_no_helix_cut** | 2 | PyRosetta version of **-splice_ncontact_no_helix_cut** |
| **--postfilt_splice_nhelix_contacted_cut** | 3 | PyRosetta version of **--splice_nhelix_contacted_cut** |

**WORMS architecture definition:**

The worms architecture can be definition in two different methods, either as an *.config file or as command line options. Described immediately below is the *.config file syntax.

```
['C3_N',orient(None,'N')),('Het:CN',orient('C','N')),('C2_C',orient('C',None)]
```

The first line of the *.config file defines the connections between all the segments, and which building blocks are allowed in each segment, as described in the main text. Marked in red is a single ‘segment’ of the worm. The first field is the ‘name’, ‘class’ or ‘type’ of the building block(s) that are desired in that segment (see database syntax for more information). The next field is ‘orient(x,y)’, which defines which connections are to be used. The termini assigned here will limit the search in the building block database to those who have that termini available. On the first and last segments, the notation ‘None’ is used to signify that there are no additional connections on that side. The segments in the center need two assignments to which termini are to be connected -- this can be ‘C’ or ‘N’, depending on what is available. In the case of a monomer, a single ‘C’ and a single ‘N’ is available. For a hetero-dimer, however, ‘N’,’N’ or ‘C’,’C’ assignments are possible. Keep in mind that a ‘N’ must connect to a ‘C’ in the next segment, and vice versa.

At time of publication, the WORMS software supports the following architectures:

```
Cyclic(symmetry=1)
D2(c2=0, c2b=-1)
D3(c3=0, c2=-1)
D4(c4=0, c2=-1)
D5(c5=0, c2=-1)
D6(c6=0, c2=-1)
Icosahedral(c5=None, c3=None, c2=None)
Octahedral(c4=None, c3=None, c2=None)
Tetrahedral(c3=None, c3b=None, c2=None)
```

The variable values listed here are the default values; they can be changed to what the user requires. For ‘Cyclic’, the symmetry variable determines what the overall oligomeric state is. For example, ‘symmetry=3’ will generate a C3 architecture. For the remaining architectures, the variables are to assign the terminal segments to their respective symmetry axis. In the ‘D3’ case, the C3 component is assigned to the ‘0th’ segment, which is the first segment listed above. The C2 component is assigned to the ‘-1th’ segment, which is the last segment listed above.

To use the command line format, the following syntax is used, a D2 architecture is shown as an example:

```
--geometry D2(c2a=0, c2b=-1)
--bbconn
      _C C2_C
      NC Monomer
      N_ C2_N
```

The major syntax difference is that instead of ‘None’, a single underscore ‘_’ is used in place for the first and last segment connections.

**WORMS database syntax and example entries:**

```
[
{"file": "/path/to/pdb/file1.pdb",
 "name": "symmetric_cyclic_example_0001" ,
 "class": ["C3_C"],
 "type": "example_C3_C" ,
 "base": "base_scaffold" ,
 "components": ["component1","component2"],
 "validated": false,
 "protocol": "made_by_example_protocol",
 "connections": [
      {"chain": 1, "direction": "C", "residues":["-129:"]},
      {"chain": 1, "direction": "N", "residues":[":180"]}
 ]
},

{"file": "/path/to/pdb/file2.pdb",
 "name": "asymmetric_het_example_0001" ,
 "class": ["Het"],
 "type": "example_het_C2_C-C" ,
 "base": "base_scaffold" ,
 "components": ["component1", "component2"],
 "validated": false,
 "protocol": "made_by_example_protocol",
 "connections": [
      {"chain": 1, "direction": "C", "residues":["-129:"]},
      {"chain": 1, "direction": "N", "residues":[":150"]},
      {"chain": 2, "direction": "C", "residues":["-139:"]},
      {"chain": 2, "direction": "N", "residues":[":86"]},
  ]
}
]
```

While not all variables need to be populated (only *file*, *name*, *class*, and *connections* are required), the other variables allow the user to customize their search of building blocks during the WORMS run. The user can specify a specific *name* in the configuration which will result in that segment being populated by a single building block. Alternatively, by searching with *class* or *type*, the user can specify that segment to be any entry that contains the desired keyword. For hetero-oligomeric entries, the class keyword “Het” is used. During the configuration setup (see above), the user can specify what kind of hetero-oligomer is desired:

`'Het:CN'` – all hetero-oligomers that have at least 1 C- and 1 N-term available.

`'Het:CNX'` – only hetero-trimers, even if you do not require the 3rd terminus

`'Het:CNY'` – only hetero-dimers.

The *base* field can be used in conjunction with the `--no_duplicate_bases` option to make sure that in a single completed architecture there will not be the same base used in non-symmetrical positions. The *components*, *validated*, and *protocols* fields are strictly for filtering purposes.

The *connections* field is where the user populates *direction*, which depicts which termini are available in each chain in a given entry. In the *residues* field, the user specifies which residues are allowed to be sampled as fusion positions. The numbering follows standard python syntax, for example, `[:100]` equates to the range: “first residue to residue 100”, and [-100:] equates to the range: “last 100 residues from the end to the end”.

**WORMS sequence design:**

All outputs from WORMS were sequence-designed using RosettaScripts with rigid backbone. The residues that need to be designed can be found appended to the WORMS asymmetric unit *.pdb output. These were identified as residues which either “gained a new contact” or “lost an old contact” in the new fused WORMS context. Each chain from the WORMS output was designed separately for computational runtime purposes, under the assumption that the junction regions are not close to one another. Afterwards, all the designed chains were then combined and designed in the symmetrical context to remove residual clashing residues. Example *.xml files can be found as supplemental files.

**Small angle X-ray scattering (SAXS):**

Sample handling and SAXS experiments were performed according to previous methods[1]. Briefly, proteins were SEC-purified in 25 mM Tris pH 8.0, 150 mM NaCl and 2% glycerol. Purified proteins collected from SEC-fractions were passed through MWCO filter columns (3 or 10 kDa cut off) to concentrate the protein samples, where the passed-through solutions were used as blanks for buffer subtraction. Scattering measurements were performed at the SIBYLS 12.3.1 beamline at the Advanced Light Source. The sample-to-detector distance was 1.5 m, and the X-ray wavelength ($\lambda$) was 1.27 Å, corresponding to a scattering vector q ($q = 4\pi \sin\theta/\lambda$, where $2\theta$ is the scattering angle) range of 0.01 to 0.3 $Å^{-1}$ . A series of exposures were taken of each well, in equal subsecond time slices: 0.3-s exposures for 10 s resulting in 32 frames per sample. Data were collected for two different concentrations for each sample: ‘low’ concentration samples ranged at 1-3 mg/ml and ‘high’ concentration samples at 2-6 mg/ml. Data was processed using the SAXS FrameSlice online server and analysed using the ScÅtter software package[2]. Experimental scattering profiles to design models were compared using the FoXS online server[3].

**X-ray crystallography:**

**X-ray crystallography Crystallization** All crystallization trials were carried out at 20°C in 96-well format using the sitting-drop method. Crystal trays were set up using Mosquito LCP by SPTLabtech. Drop volumes ranged from 200 to 400 nl and contained protein to crystallization solution in ratios of 1:1, 2:1 and 1:2. Diffraction quality crystals appeared in 0.2M Sodium chloride, 0.1M Sodium/Potassium phosphate pH 6.2 and 50% PEG200 (JCSG+ D3) for C3_HDock-1069; 0.2 M Lithium sulfate, 0.1M Na-acetate pH 4.5 and 2.5 M NaCl for C3_nat_HF-0005; 0.2 M $MgCl_2$, 0.1 TrisCl pH 8.5, 10% Glycerol and 25% (v/v) 1,2-Propanediol for C3_HF_Wm-0024A; and 0.1M MES pH 5.0, 20% MPD plus an additional 20% MPD as a cryoprotectant for C3_Crn-05. Crystals were subsequently harvested in a cryo-loop and flash frozen directly in liquid nitrogen for synchrotron data collection.

**X-ray crystallography Data Collection** Data collection from crystal of C3_nat_HF-0005 was performed with synchrotron radiation at the Advanced Photon Source (APS), 24ID-E. Crystals belonged to space group R 3 :H with cell dimensions a = b = 101.97 Å, and c = 78.44 Å, α = β = 90° and γ = 120°. Data collection from the crystal of C3_HF_Wm-0024A was performed with synchrotron radiation at the Advanced Light Source (ALS), 8.2.2. Crystals belonged to space group $P4_32_12$ with cell dimensions a = b = 166.77 Å, and c = 223.51 Å, α = β = γ = 90°. X-ray intensities and data reduction were evaluated and integrated using XDS[4] and merged/scaled using Pointless/Aimless in the CCP4 program suite[5].

**Structure determination and refinement** Starting phases were obtained by molecular replacement using Phaser[6] using the designed model for the structures. Following molecular replacement, the models were improved using Phenix autobuild[7]; efforts were made to reduce model bias by setting rebuild-in-place to false, and using simulated annealing and prime-and-switch phasing. Structures were refined in Phenix. Model building was performed using COOT[8]. The final model was evaluated using MolProbity[9]. Data collection and refinement statistics are recorded in Table S1.

**Data deposition** The crystallography, atomic coordinates, and structure factors reported in this paper have been deposited in the Protein Data Bank (PDB), http://www.rcsb.org/ with accession codes 6XH5, 6XI6, 6XNS and 6XT4.

**Electron microscopy: cyclic structures (C4, C5 and C6)**

**Negative stain EM grid preparation, data collection, and data processing** Proteins were diluted to 20 μg/ml with TBS, then immediately applied to freshly glow-discharged Formvar/carbon 400 mesh copper grids (Ted Pella catalog #01754-F). After incubation for 45s, excess protein solution was removed by blotting from the side with filter paper, then grids were inverted onto two successive drops of sample buffer followed by three to five successive drops of 2% uranyl formate, with excess solution removed by blotting after each application. The final stain applied was incubated for 15s before blotting. Air-dried grids were imaged using a FEI Talos L120C TEM equipped with a 4K × 4K Gatan OneView camera, at a nominal magnification of 73,000x and pixel size of 2.0 Å. Micrographs were imported to Relion 3.1[10] and/or cryoSPARC v2[11] and, after picking using automated protocols in each program, particles were subjected to 2D classification. Design model projections were generated using EMAN2[12] and Relion, and projections were aligned with experimental 2D class averages using Sparx[13].

**Cryo-EM grid preparation and data collection** 3.5 µL of C4_nat_HF-7900 at a concentration of 1 mg/ml was applied to 400 mesh copper Quantifoil holey carbon grids 1.2/1.3 coated with graphene oxide (catalog #GOQ400R1213Cu, Electron Microscopy Sciences). C5_HF-3921 and C5_HF-0019 were diluted with TBS to final concentrations of 0.75 mg/ml and 0.45 mg/ml, respectively, immediately before applying to glow-discharged 400 mesh copper Quantifoil holey carbon grids 1.2/1.3 (3.5 µL of C5_HF-3921 and 3.0 µL of C5_HF-0019). All grids were plunge-frozen using a Vitrobot Mark IV. Grids were pre-screened on a Talos Arctica microscope operated at 200 kV with a Gatan K3 camera (NYU) and C4_nat_HF-7900 movies were collected with this setup. C5_HF-3921 movies were acquired on a Titan Krios microscope ("Krios 3") operated at 300 kV with Gatan K3 camera and located at the New York Structural Biology Center. To address preferred orientation of particles, C5_HF-3921 movies were acquired at both 0° and 35° tilt angles, and for tilted movies a 4s pre-exposure wait time was added. Data acquisition was controlled via Leginon[14] and pre-processing was performed with Appion[15]. Data collection parameters are shown in Supplementary Table S2.

**Cryo-EM data processing** Detailed processing workflows are shown in Figure S7 and S9. Movies were motion-corrected and dose-weighted using MotionCor2[16] within Leginon/Appion, then imported to cryoSPARC v2 for CTF estimation, particle picking, 2D classification, and *ab initio* 3D reconstruction. For C4_nat_HF-7900, particles picked "on-the-fly" with Warp[17] were imported to cryoSPARC for 2D classification to generate particles to use as templates for template-based picking. Multiple rounds of 2D classification and manual curation were used to generate a set of particles to use as a training set for Topaz[18]. Topaz-picked particles were then used for further 2D/3D classification and 3D refinement in cryoSPARC. For C5_HF-3921, images collected at both 0° and 35° were processed together following patch CTF estimation for 2D classification, *ab initio* 3D reconstruction, and initial 3D refinement. The best C5_HF-3921 map resulted from 3D refinement of data collected at a 35° tilt angle only. For C4_nat_HF-7900, after initial processing in cryoSPARC, particles picked by Topaz were imported to Relion 3.1 for further 2D/3D classification and 3D refinement. 3D refinements were performed both with and without symmetry imposed. For C4_nat_HF-7900, imposing C4 symmetry yielded the highest quality map, whereas for C5_HF-3921 a C1 map had higher overall quality despite lower nominal resolution (due to artifacts introduced by imposing C5 symmetry). Overall map resolutions were estimated using the gold-standard Fourier Shell Correlation criterion (FSC=0.143) within Relion (C4_nat_HF-7900) or cryoSPARC (C5_HF-3921) and 3D FSC were calculated using the "Remote 3DFSC Processing Server" (https://3dfsc.salk.edu/)[19]. Soft masks were provided for estimation of local resolution of C4_nat_HF-7900 and C5_HF-3921 maps using implementations within Relion and cryoSPARC, respectively.

**C4_nat_HF-7900 model building and refinement** The *ab initio* coordinates of the C4_nat_HF-7900 design were used as the starting model. Four C4_nat_HF-7900 protomers were first individually docked into the cryo-EM map as rigid bodies using Chimera[20], then refined using iterative rounds of refinement with real_space_refine in PHENIX[7] followed by manual model adjustment in COOT[21,22]. Each of the four chains in the tetramer was divided into 2 rigid bodies (residues 1-65 and 66-295; corresponding to the HB and DHR, respectively). Rigid body and ADP refinement were performed, with secondary structure, non-crystallographic symmetry, Ramachandran, and rotamer restraints enabled. The model was then analyzed using COOT and residues 261 - 295 were removed due to weak density in this region of the cryo-EM map. Since the C-terminus of C4_nat_HF-7900 is >95% identical to a previously characterized DHR (PDB ID: 5cwp[23]), secondary structure restraints for residues 71 - 260 were based on the 5cwp structural model. After multiple iterations of real_space_refine and manual model adjustment, all helices except for the two C-terminal helices in the model (residues 212 - 260) were well-placed within the cryo-EM map density. Inspection of the model and map showed ambiguity in the position of residues 210 - 213 due to low local resolution and discontinuous density in this region. This loop and the following two C-terminal helices were shifted relative to their position in the 5cwp structure, possibly as a result of incorrect Thr210 - Pro213 loop placement. To determine whether this shift reflected a true difference between the C4_nat_HF-7900 DHR and 5cwp structures, we used the 5cwp structural model to drive placement of these helices as follows: 5cwp was aligned to residues 101 - 260 of the working C4_nat_HF-7900 model (excluding the N-terminal DHR helix in case of distortions introduced from fusion to the HB) and a hybrid model was created by joining residues 1 - 208 of C4_nat_HF-7900 to residues 140 - 191 of 5cwp using Chimera. The single amino acid difference between C4_nat_HF-7900 and 5cwp in the grafted C-terminus was mutated to restore the C4_nat_HF-7900 design sequence, and this model was subjected to additional rounds of PHENIX real_space_refine and manual refinement in COOT. After refinement, the backbone of the Thr210 - Pro213 loop and C-terminal helices remained in position, leading to close alignment of C4_nat_HF-7900 residues 101 - 260 with 5cwp and a better fit of the two C-terminal helices to the cryo-EM map density.

**Electron microscopy: higher order structures (crowns, dihedrals, and the point group cages):**

**Negative-stain electron microscopy (NS-EM)** Negative-stained sample grids for transmission electron microscopy were prepared using either Nano-W or Uranyl Formate (Nanoprobes) at a sample concentration of 0.01-0.005 mg/mL using manufacturer's standard operating procedure. Stained grids were screened using FEI Morgagni transmission electron microscope operating at 100 kV. For 2D averaging, images were collected in a Tecnai T12 electron microscope using Leginon image collection software. The parameters of the contrast transfer function (CTF) were estimated using CTFFIND4. All particles were picked in a reference-free manner using

DoG Picker. Reference-free 2D classification was used to select homogeneous subsets of particles using CryoSPARC. The selected particles were subsequently subjected to ab initio 3D reconstructions and Homogenous 3D refinement using CryoSPARC.

**Cryo-electron microscopy** 3 µL of 1 mg ml -1 of C5_Crn_HF_12_26 was loaded onto a freshly glow-discharged (30 s at 20 mA) 1.2/1.3 UltraFoil grid (300 mesh) prior to plunge freezing using a vitrobot Mark IV (ThermoFisher Scientific) using a blot force of 0 and 6 second blot time at 100% humidity and 25°C. Data was acquired using an FEI Glacios transmission electron microscope operated at 200 kV and equipped with a Gatan K2 Summit direct detector. Automated data collection was carried out using Leginon at a nominal magnification of 36,000x with a pixel size of 1.16 Å. The dose rate was adjusted to 8 counts/pixel/s, and each movie was acquired in counting mode fractionated in 50 frames of 200 ms. 1,709 micrographs were collected with a defocus range between -1.0 and -3.5 µm. Movie frame alignment, estimation of the microscope contrast-transfer function parameters, particle picking, and extraction were carried out using Warp. Reference-free 2D classification was used to select homogeneous subsets of particles using CryoSPARC. The selected particles were subsequently subjected to ab initio 3D reconstructions and 3D refinements using CryoSPARC.

3 µL of 1 mg ml -1 of I32_Wm-42 was loaded onto a freshly glow-discharged (30 s at 20 mA) 2.2um c-flat grid prior to plunge freezing using a vitrobot Mark IV (ThermoFisher Scientific) using a blot force of 0 and 6 second blot time at 100% humidity and 25°C. Data were acquired using the an FEI Glacios transmission electron microscope operated at 200 kV and equipped with a Gatan K2 Summit direct detector.

Automated data collection was carried out using Leginon at a nominal magnification of 36,000x with a pixel size of 1.16 Å. 618 micrographs were collected with a defocus range between -1.2 μm and -3.5 μm. Movie frame alignment and estimation of the microscope contrast-transfer function parameters were carried out using Warp. 500 particles were picked initially and 2D classifications were performed in cisTEM. Eleven representative 2D class averaged images were selected as references for automatic particle picking. 2D classifications were performed in RELION 3.0. The selected particles were subsequently subjected to ab initio 3D reconstructions using CryoSPARC. 3D classification and 3D refinements were performed using RELION 3.0.

### 2) SUPPLEMENTARY FIGURES

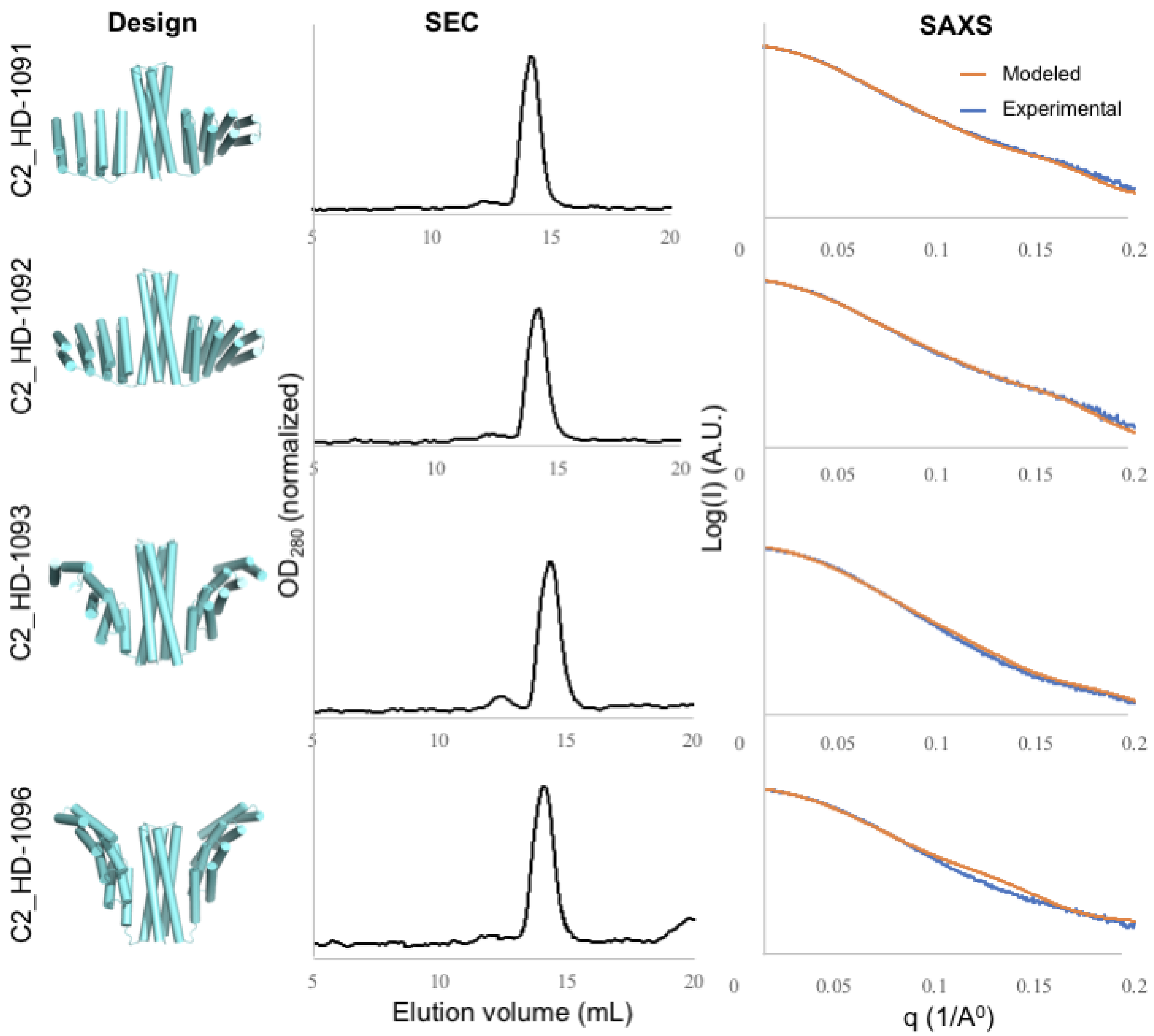

**Figure S1**: **SEC and SAXS characterizations of C2 symmetric oligomers which were designed using the 'HelixDock' protocol**. The *left* panel shows the designed models; the *middle* shows the SEC curves (Superdex 200), and the *right* shows the SAXS fitting comparison between the designed model (orange) and the experimental data (blue).

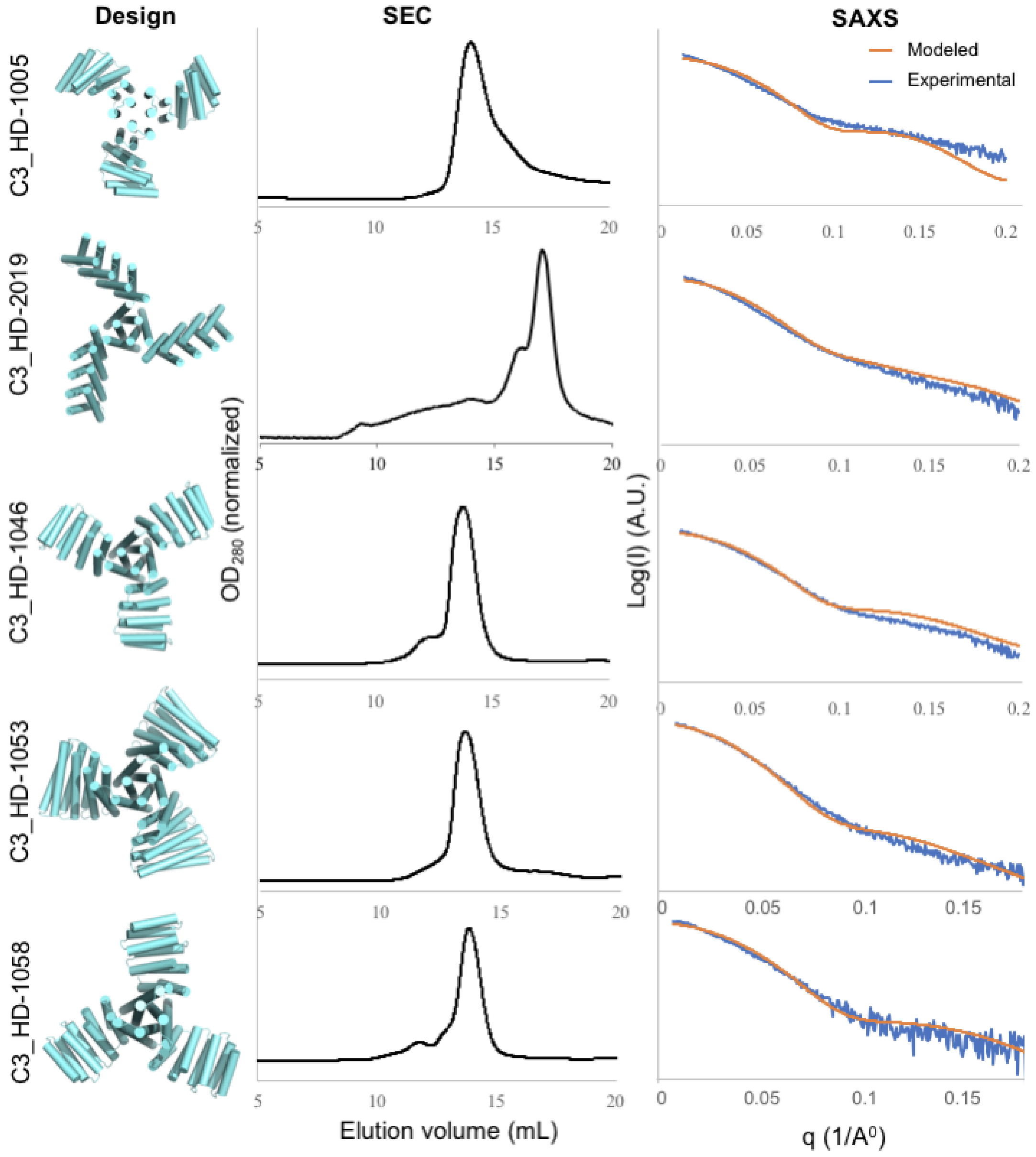

**Figure S2A**: **SEC and SAXS characterizations of C3 symmetric oligomers which were designed using the 'HelixDock' protocol**. The *left* panel shows the designed models; the *middle* shows the SEC curves (Superdex 200), and the *right* shows the SAXS fitting comparison between the designed model (orange) and the experimental data (blue).

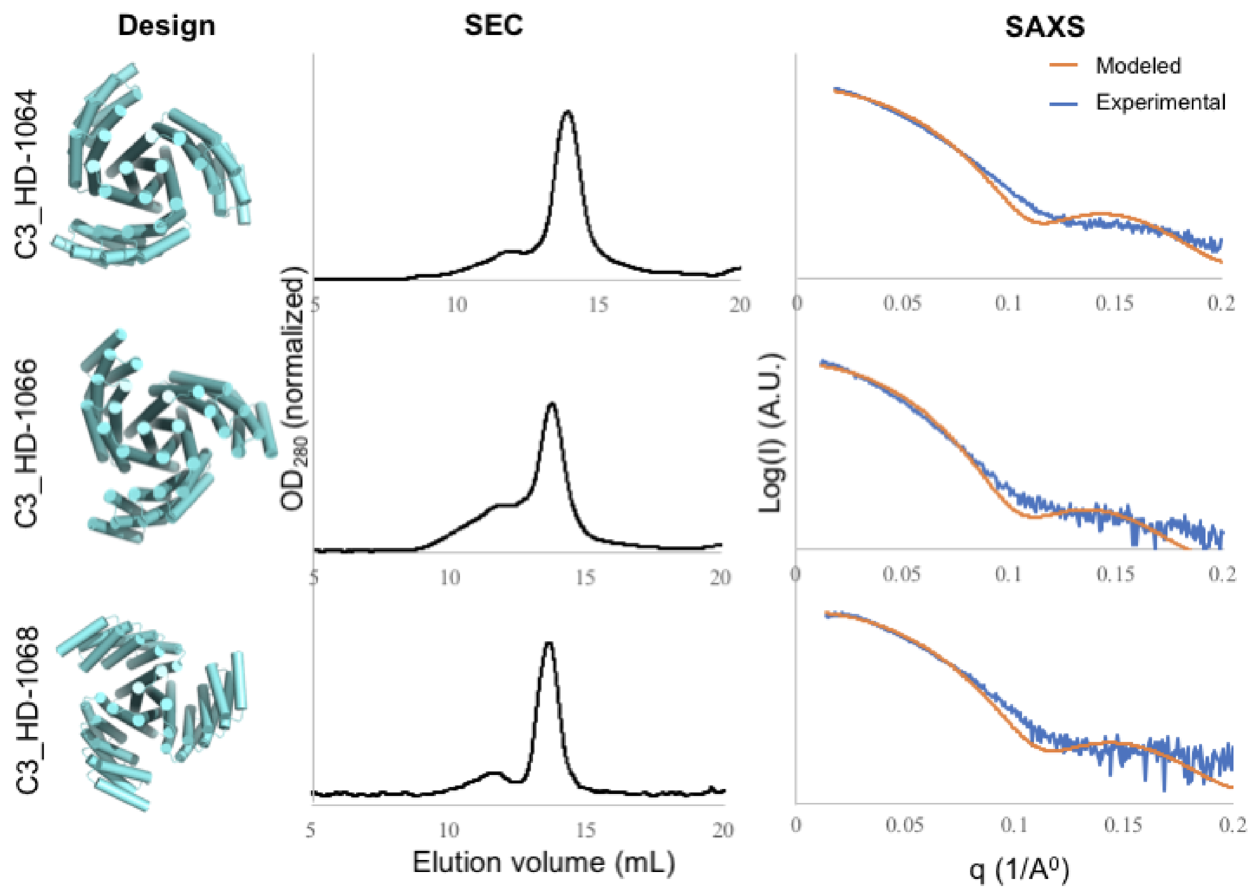

**Figure S2B**: **SEC and SAXS characterizations of C3 symmetric oligomers which were designed using the 'HelixDock' protocol**. The *left* panel shows the designed models; the *middle* shows the SEC curves (Superdex 200), and the *right* shows the SAXS fitting comparison between the designed model (orange) and the experimental data (blue).

397 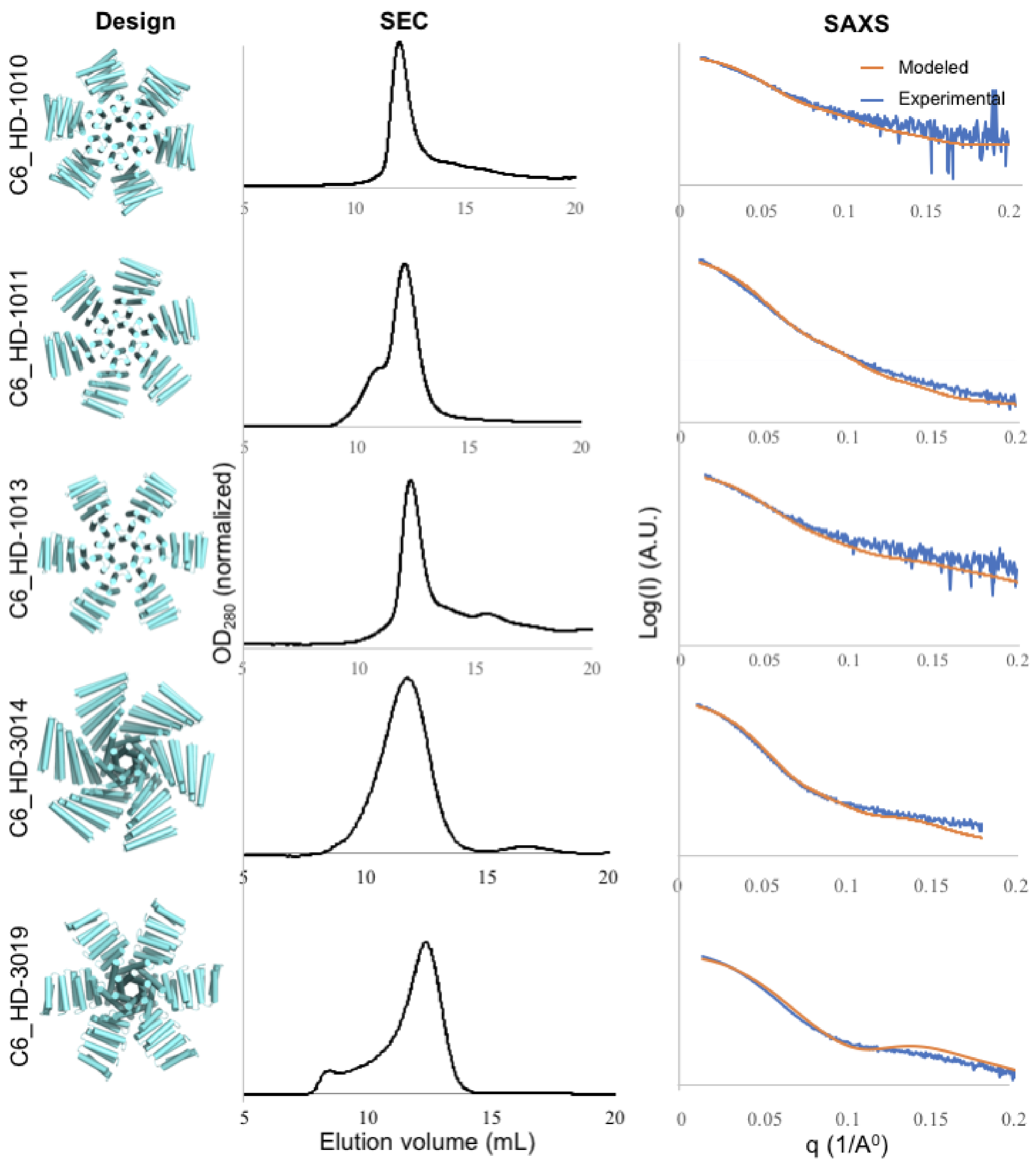

398 **Figure S3**: **SEC and SAXS characterizations of C6 symmetric oligomers which**
399 **were designed using the 'HelixDock' protocol**. The *left* panel shows the designed
400 models; the *middle* shows the SEC curves (Superdex 200), and the *right* shows the
401 SAXS fitting comparison between the designed model (orange) and the experimental
402 data (blue).

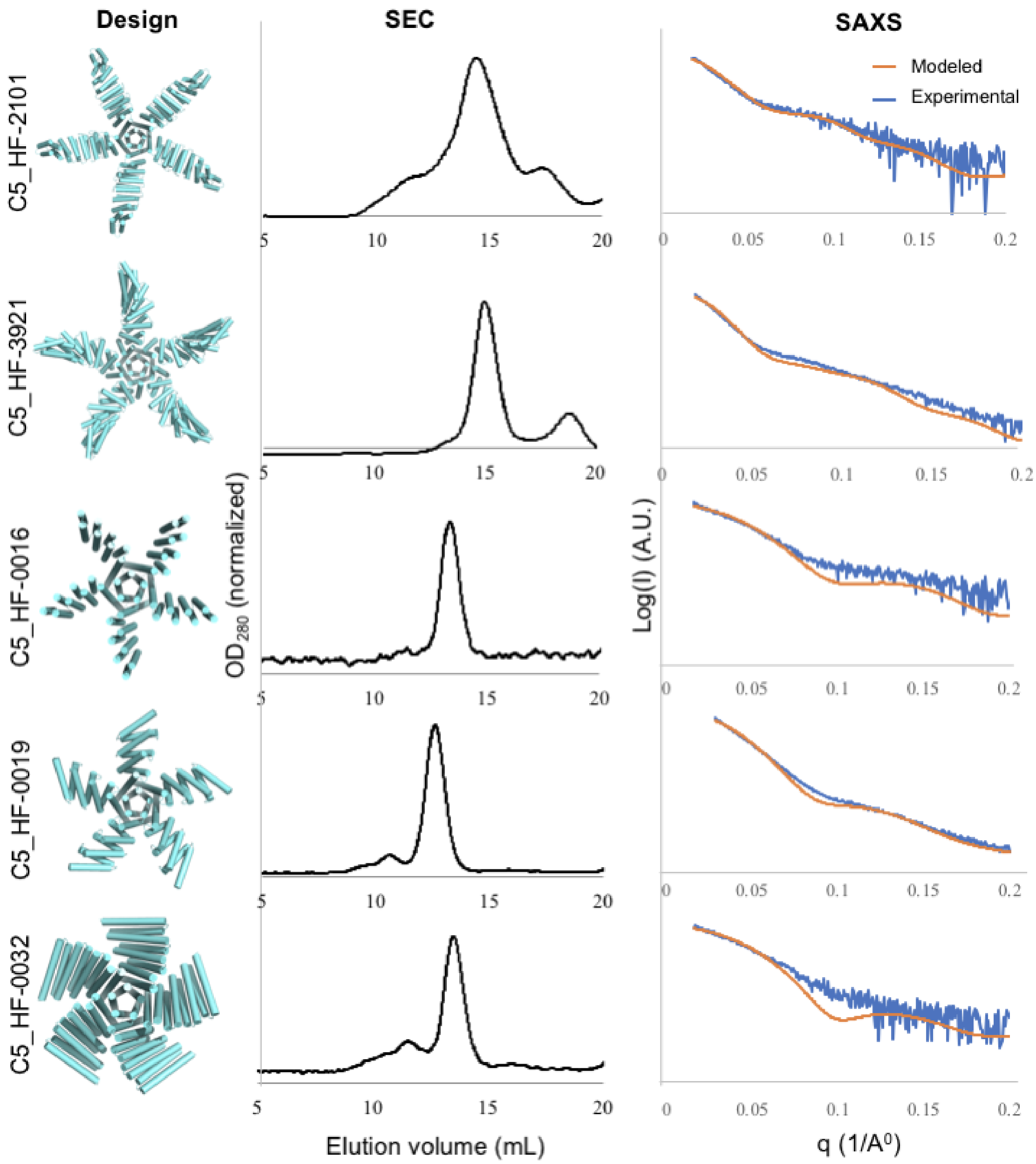

**Figure S4**: **SEC and SAXS characterizations of C5 symmetric oligomers which were designed using the 'HelixFuse' protocol**. The *left* panel shows the designed models; the *middle* shows the SEC curves (C5_HF-2101 and C5_HF-3921 by Superose 6, remaining by Superdex 200), and the *right* shows the SAXS fitting comparison between the designed model (orange) and the experimental data (blue).

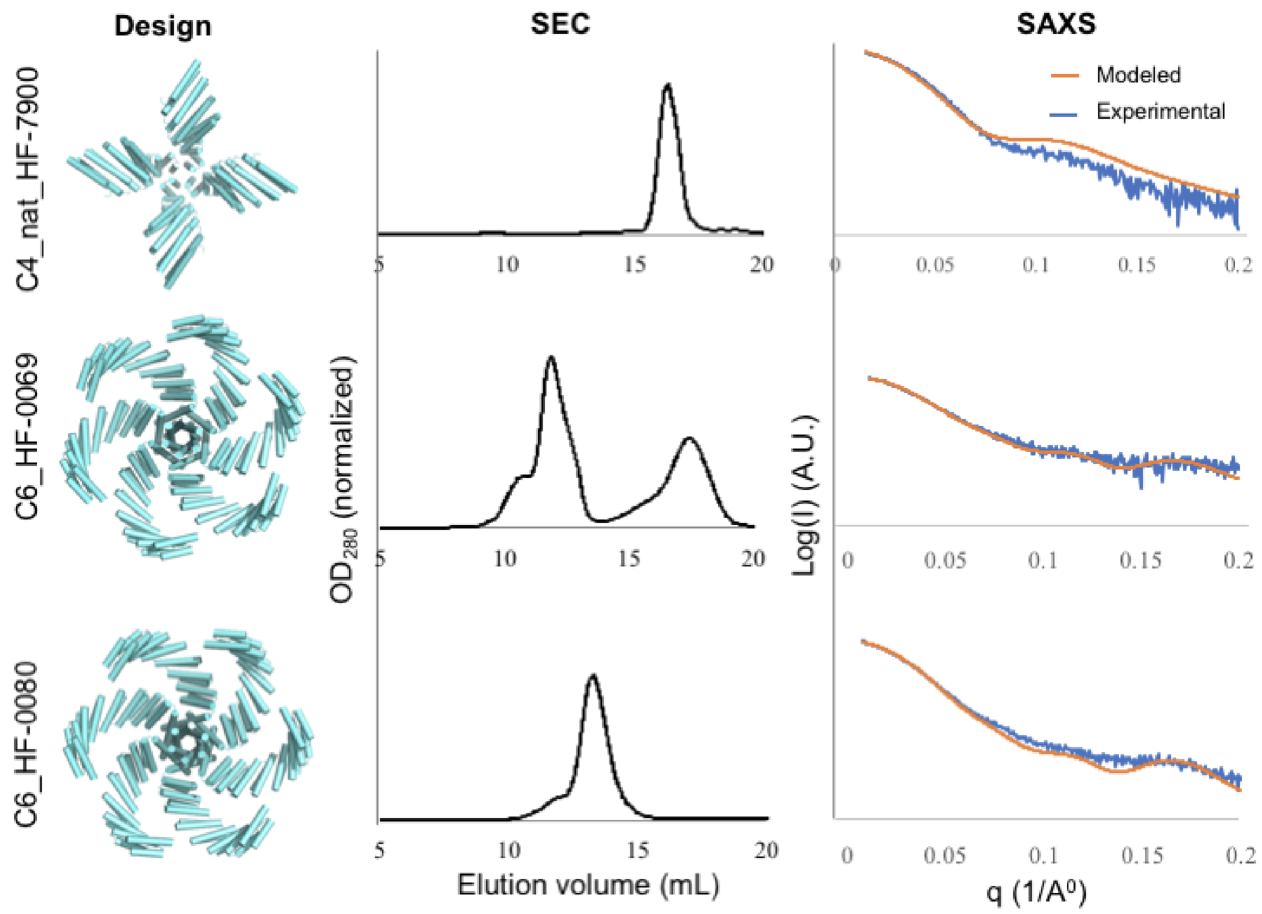

**Figure S5**: **SEC and SAXS characterizations of C4 and C6 symmetric oligomers which were designed using the 'HelixFuse' protocol**. The *left* panel shows the designed models; the *middle* shows the SEC curves (C4_nat_HF-7900 by Superose 6, C6_HF-0069 and C6_HF-0080 by Superdex 200), and the *right* shows the SAXS fitting comparison between the designed model (orange) and the experimental data (blue).

417

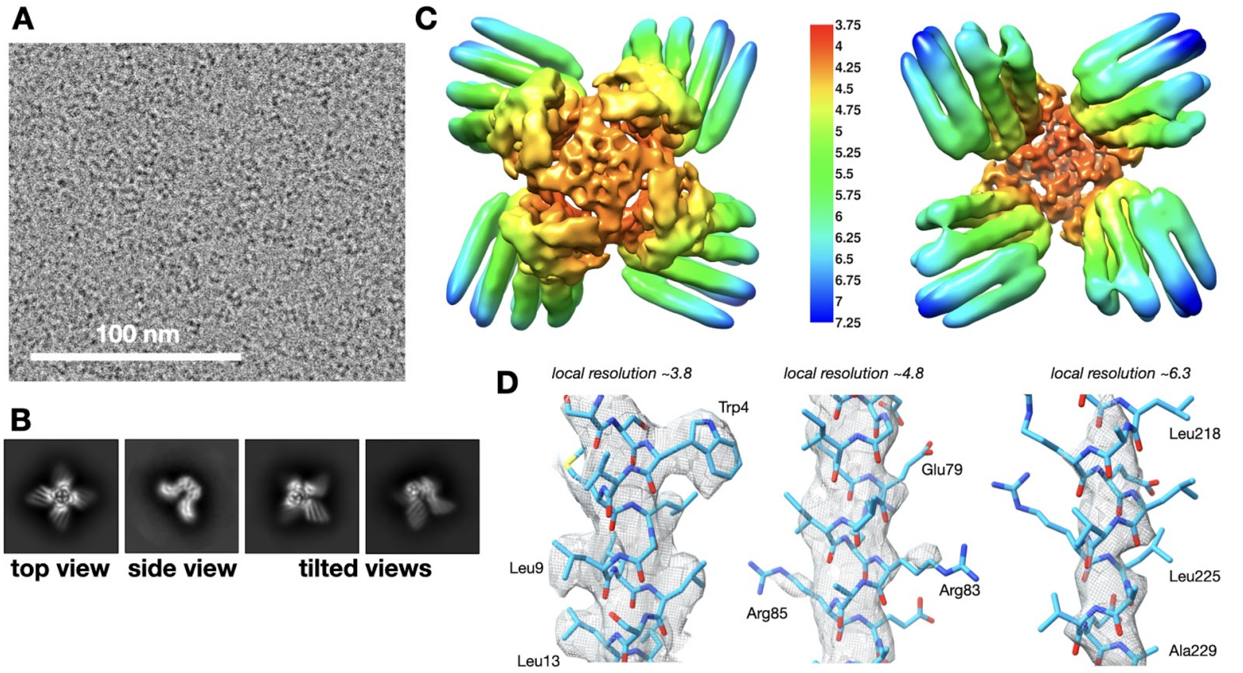

418 **Figure S6: Cryo-EM data and reconstruction for C4_nat_HF-7900.** (A)
419 Representative motion-corrected micrograph. (B) Representative 2D class averages.
420 (C) Locally-filtered cryo-EM map colored by local resolution. (D) Fit of cryo-EM structure
421 (sticks) to density (mesh) in areas of high, intermediate, and low local resolution.

422

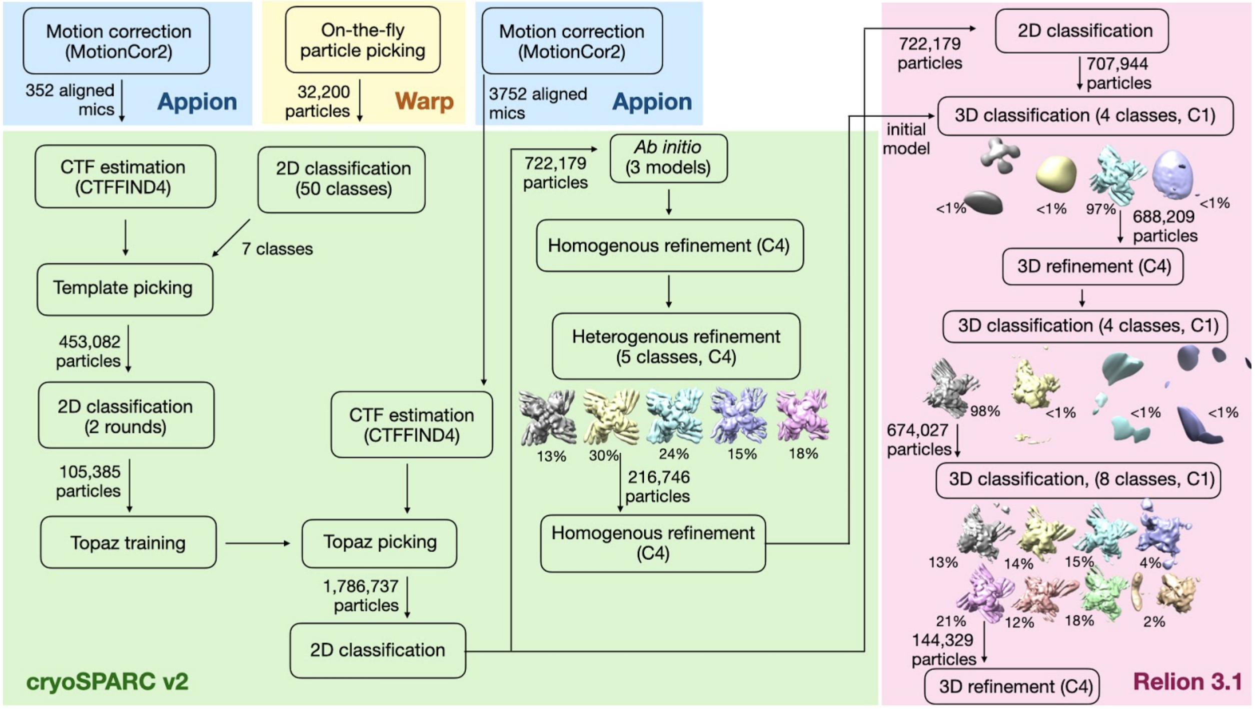

423 **Figure S7: Cryo-EM data processing workflow for C4_nat_HF-7900.**

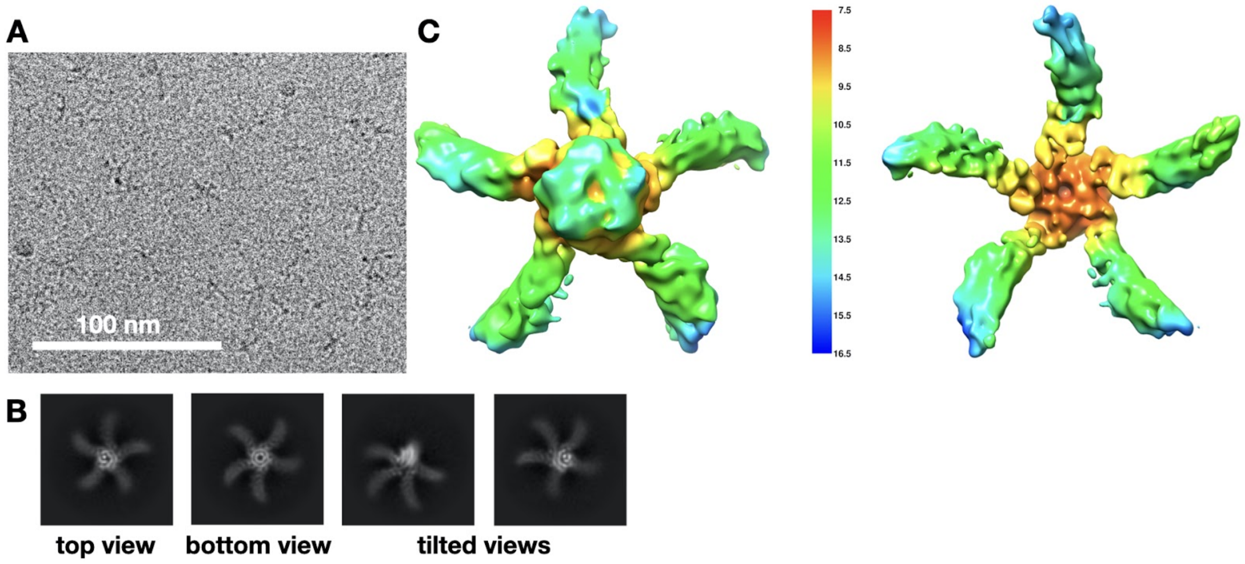

**Figure S8: Cryo-EM data for C5_HF-3921.** (A) Representative motion-corrected micrograph. (B) Representative 2D class averages. (C) Cryo-EM map colored by local resolution.

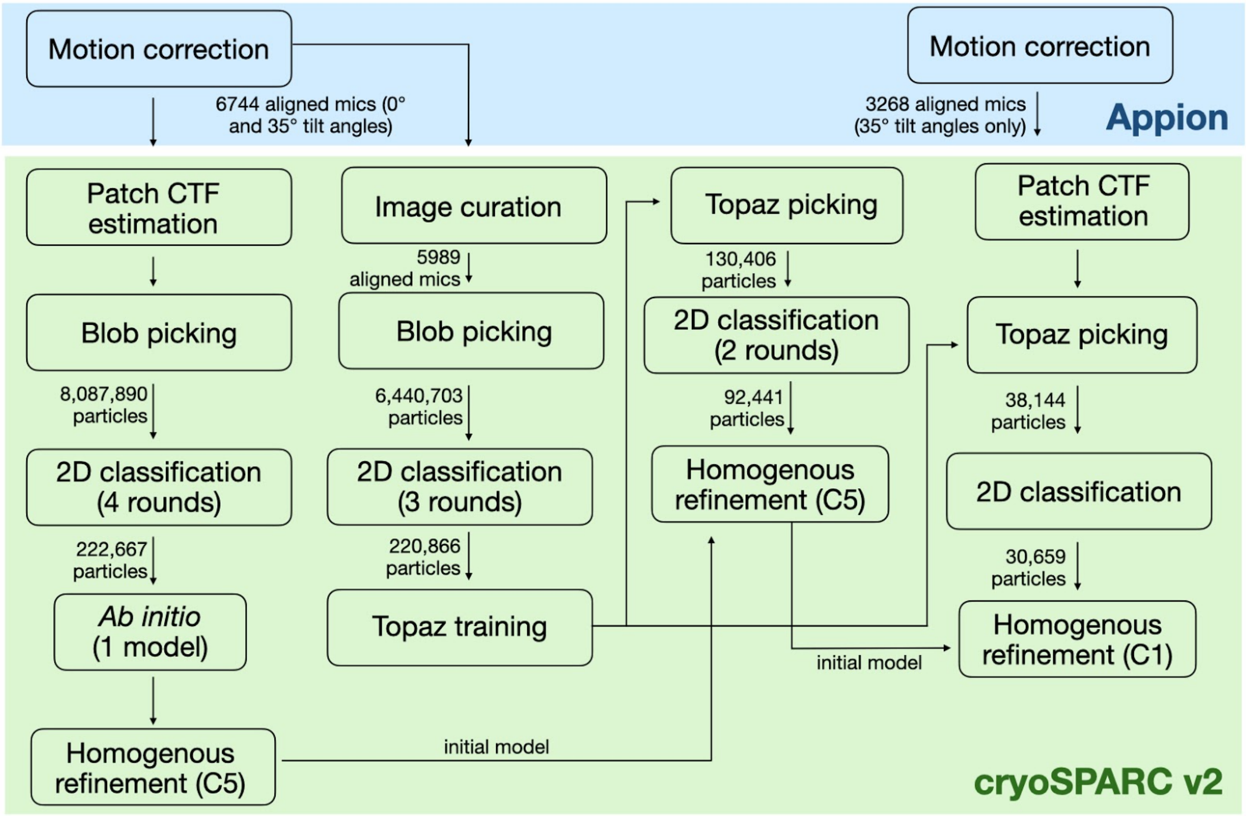

**Figure S9: Cryo-EM data processing workflow of C5_HF-3921.**

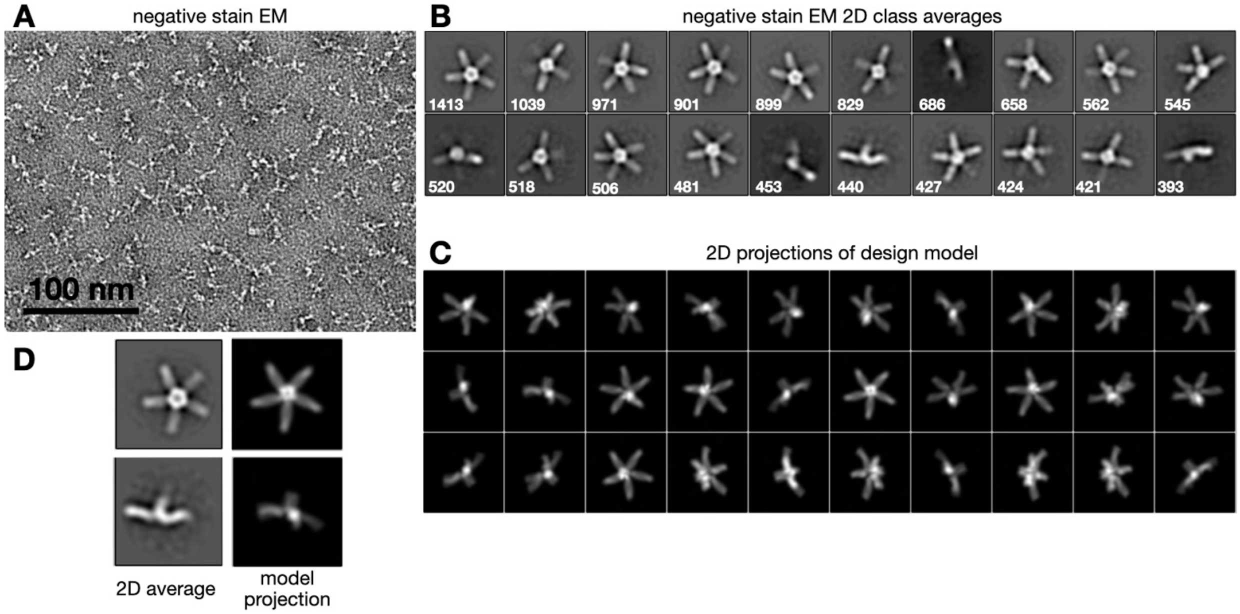

**Figure S10: Negative stain EM data for C5_HF-2101**. (A) Representative micrograph. (B) Most populated 2D class averages; numbers on each class image indicate the number of particles in that class. (C) 2D projections of a 20 Å-filtered volume generated from the atomic coordinates of the design model. (D) Selected 2D class averages shown alongside matching model projections.

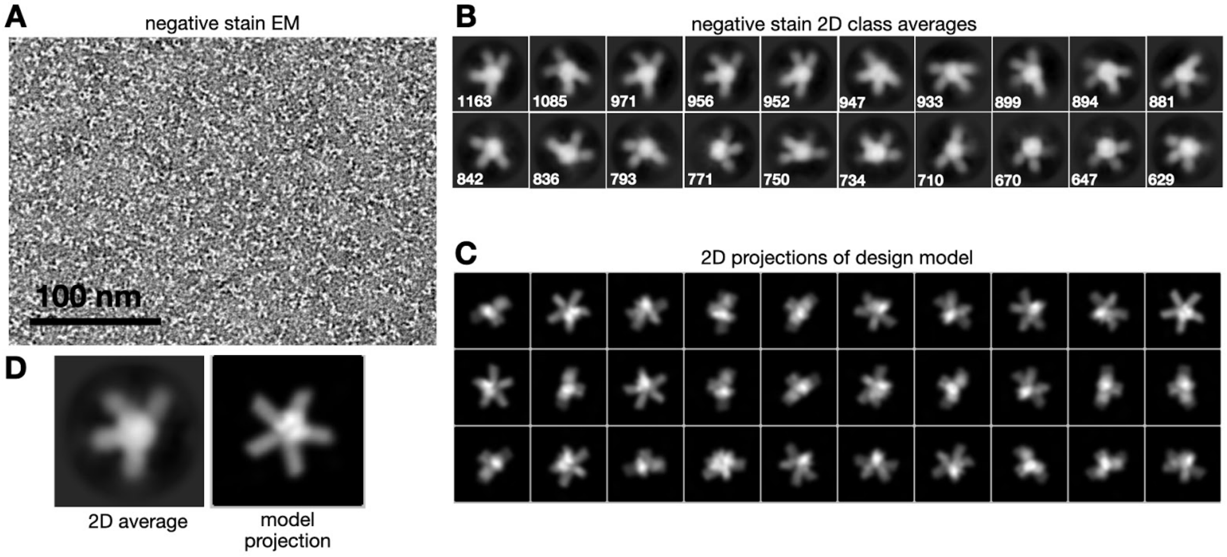

**Figure S11. Negative stain EM data for C5_HF-0007.** (A) Representative micrograph. (B) Most populated 2D class averages; numbers on each class image indicate the number of particles in that class. (C) 2D projections of a 20 Å-filtered volume generated from the atomic coordinates of the design model. (D) Selected 2D class average shown alongside matching model projection.

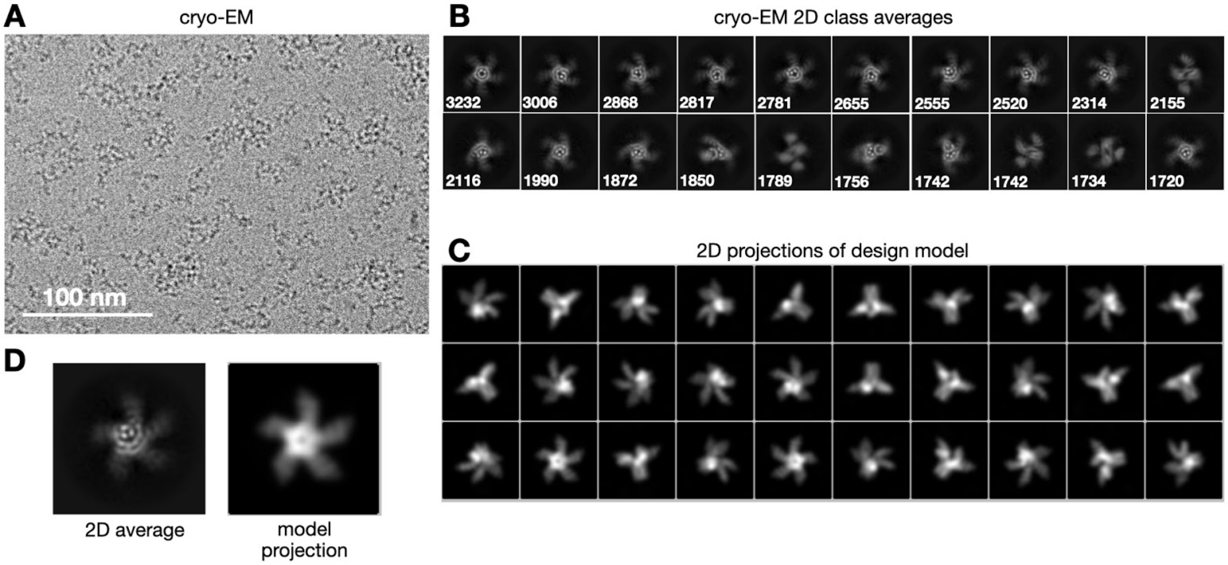

**Figure S12: Cryo-EM data for C5_HF-0019.** (A) Representative motion-corrected micrograph. (B) Most populated 2D class averages; numbers on each class image indicate the number of particles in that class. (C) 2D projections of a 15 Å-filtered volume generated from the atomic coordinates of the design model. (D) Selected 2D class average shown alongside matching model projection.

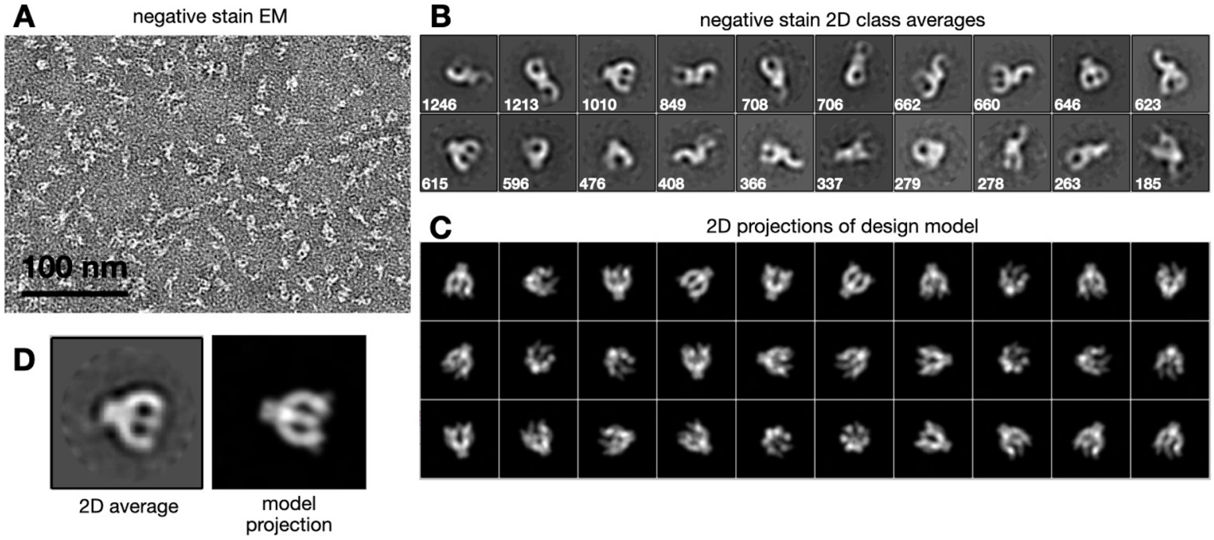

**Figure S13. Negative stain EM data for C6_HF-0075.** (A) Representative micrograph. (B) Most populated 2D class averages; numbers on each class image indicate the number of particles in that class. (C) 2D projections of a 20 Å-filtered volume generated from the atomic coordinates of the design model. (D) Selected 2D class average shown alongside matching model projection.

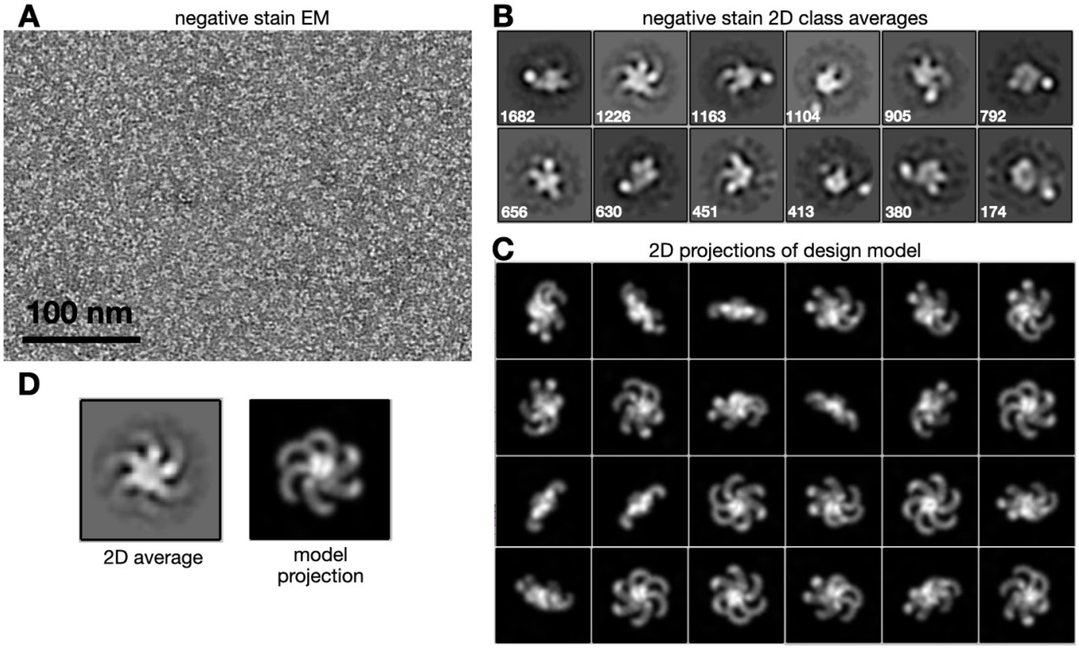

**Figure S14. Negative stain EM data for C6_HF-0080.** (A) Representative micrograph. (B) Most populated 2D class averages; numbers on each class image indicate the number of particles in that class. (C) 2D projections of a 20 Å-filtered volume generated from the atomic coordinates of the design model. (D) Selected 2D class average shown alongside matching model projection.

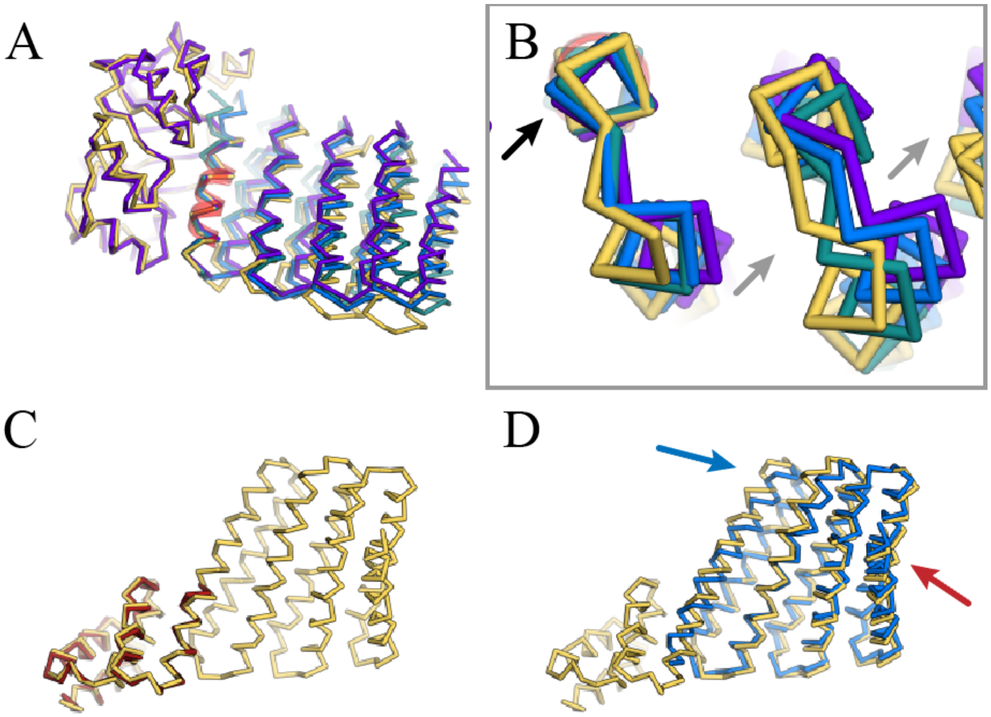

**Figure S15: Alignment of the original scaffolds to C3_nat_HF-0005's crystal structure and C4_nat_HF-7900's cryo-EM model**. Symmetric units hidden for clarity. A) C3_nat_HF-0005 design mode (purple) and crystal structure (yellow), aligned at the 1wa3 hub. DHR49 model (blue) and DHR49's original crystal structure (teal) aligned at the junction helix, highlighted in red. B) A small deviation in the loop region of the first helix (black arrow) propagates into a large deviation towards the distal portion of the protein (grey arrows). C) tpr1C4_pm3 (red) aligned to C4_nat_HF-7900's cryo-EM model (yellow). D) DHR79 (blue) aligned to C4_nat_HF-7900's cryo-EM model (yellow). While the majority of the DHR aligns well, the N-terminal helices align less well to the model regardless of the new fusion region (blue arrow). The C-terminal helix is not present in the cryo-EM map (red arrow).

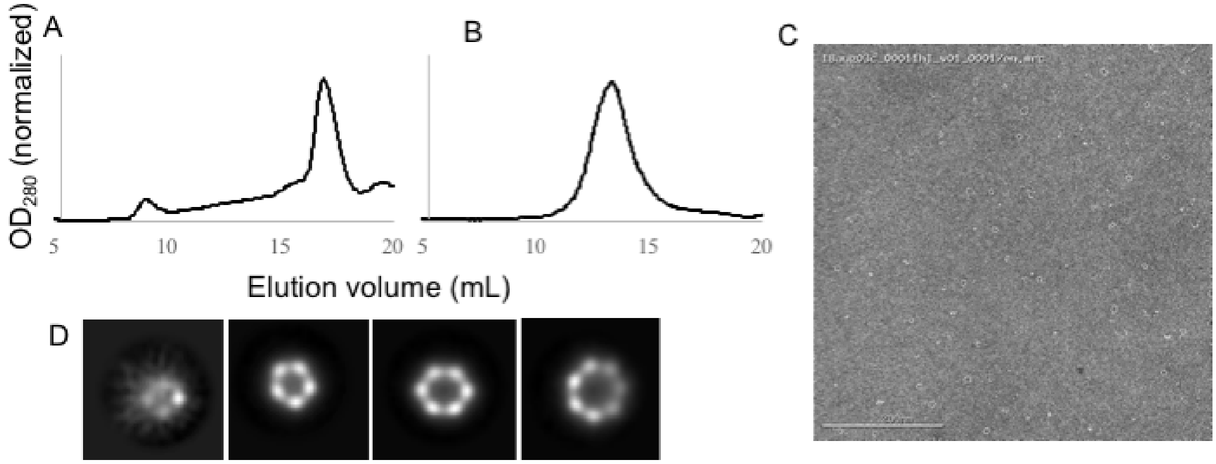

**Figure S16**: **Characterization of C3 and C5 crowns**. SEC of A) C3_Crn-05 (Superdex 200), and B) C5_Crn-07 (Superdex 200). C) C5_Crn-07 negative stain micrograph, D) C5_Crn-07 negative stain 2D average showing all alternative states.

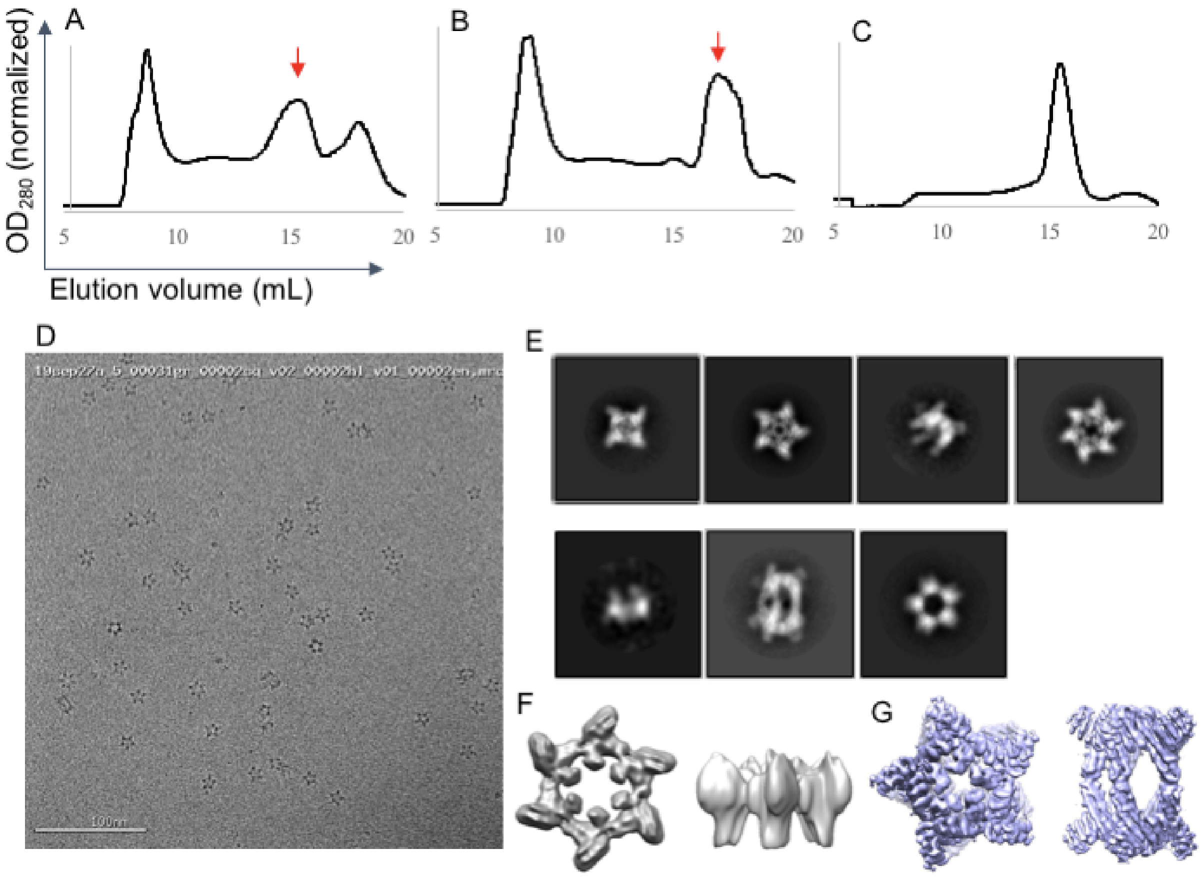

**Figure S17**: **Characterization of C5_Crn-07 with extended arms**. SEC of A) C5_Crn_HF-12 (Superose 6), B) C5_Crn_HF-26 (Superose 6), and C) C5_Crn_HF-12_26 (Superose 6). Red arrows indicate the correct elution fractions; aggregate fraction for A and B were disregarded. Cryo electron microscopy characterization of C5_Crn_HF-12_26. D) representative micrograph; E) class averages showing off-target states. Cryo-EM density maps for additional off-target states: F) C6 (*left*--top view, *right*--side view), G) D5 (*left*--top view, *right*--side view).

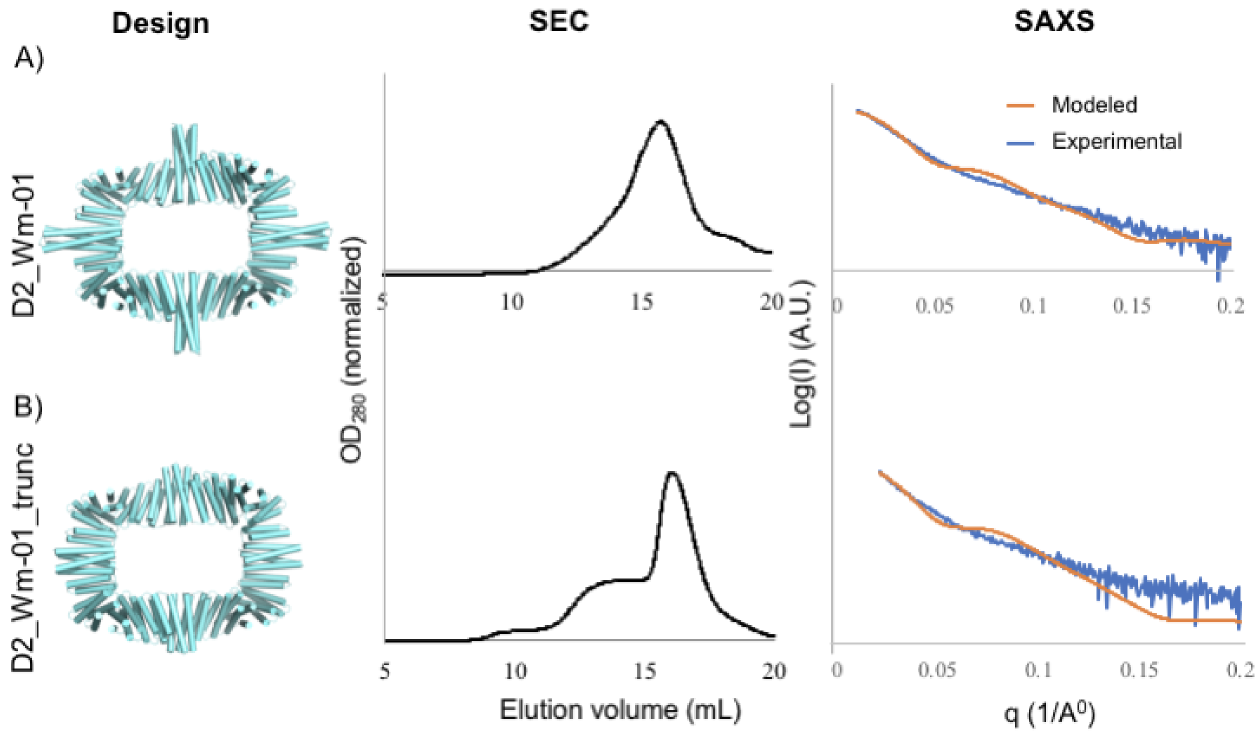

**Figure S18**: **Characterization of D2_Wm-01 (A) and D2_Wm-01_trunc (B) dihedral rings.** The *left* panel shows the designed models; the *middle* panel shows the SEC curves (Superose Increase 10/300 S6 column); and the *right* panel shows SAXS fitting curves which were compared between the designed model (orange) and the experimental data (blue).

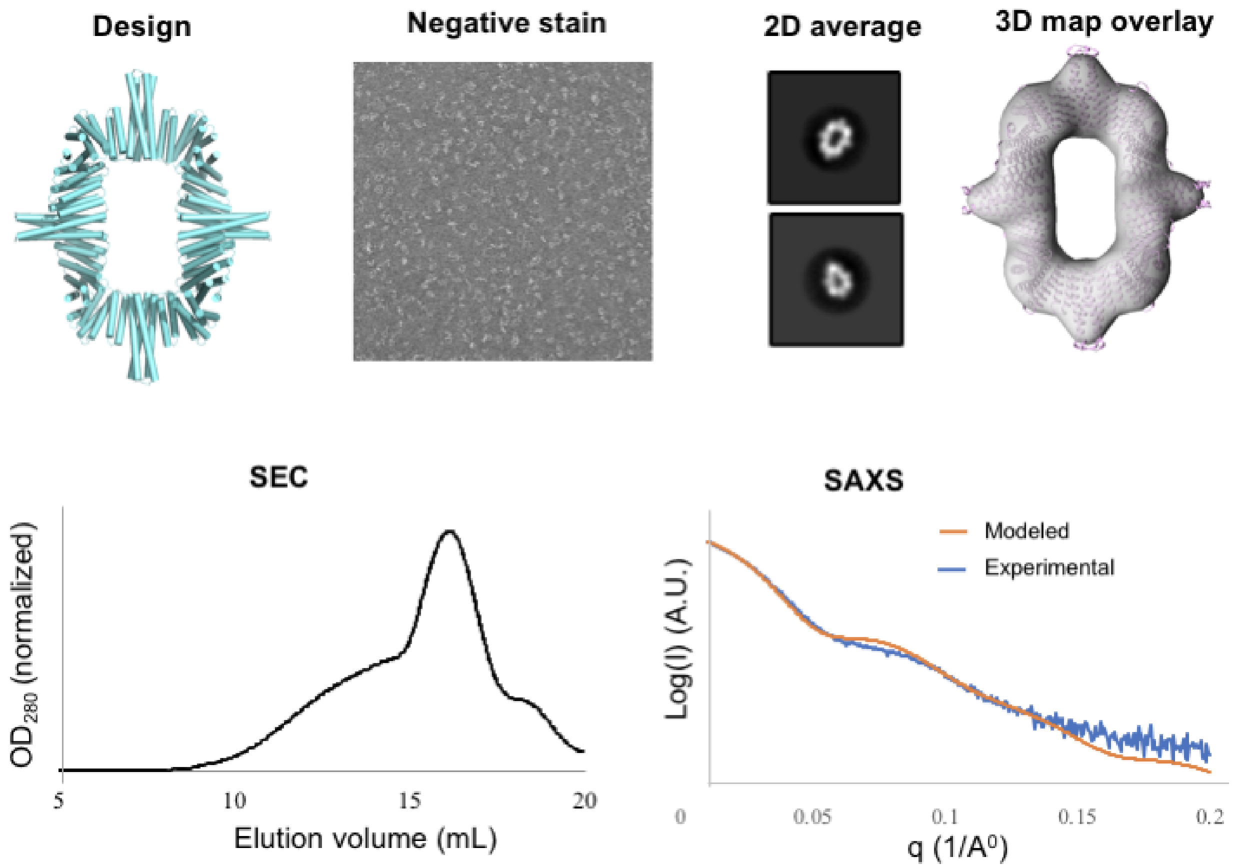

**Figure S19: Characterization of D2_Wm-02 dihedral ring**. Two-component D2_Wm-02 ring was designed using the WORMS protocol, which was then expressed and subsequently purified using SEC (Superose Increase 10/300 S6 column). Purified protein was characterized by either SAXS or NS EM. 2D average of the NS EM shows features resembling the designed model, and the 3D density map (upper right) overlays accurately with the designed model. Likewise, SAXS fitting (lower right) shows the close resemblance between the designed model (orange) and the experimental data (blue).

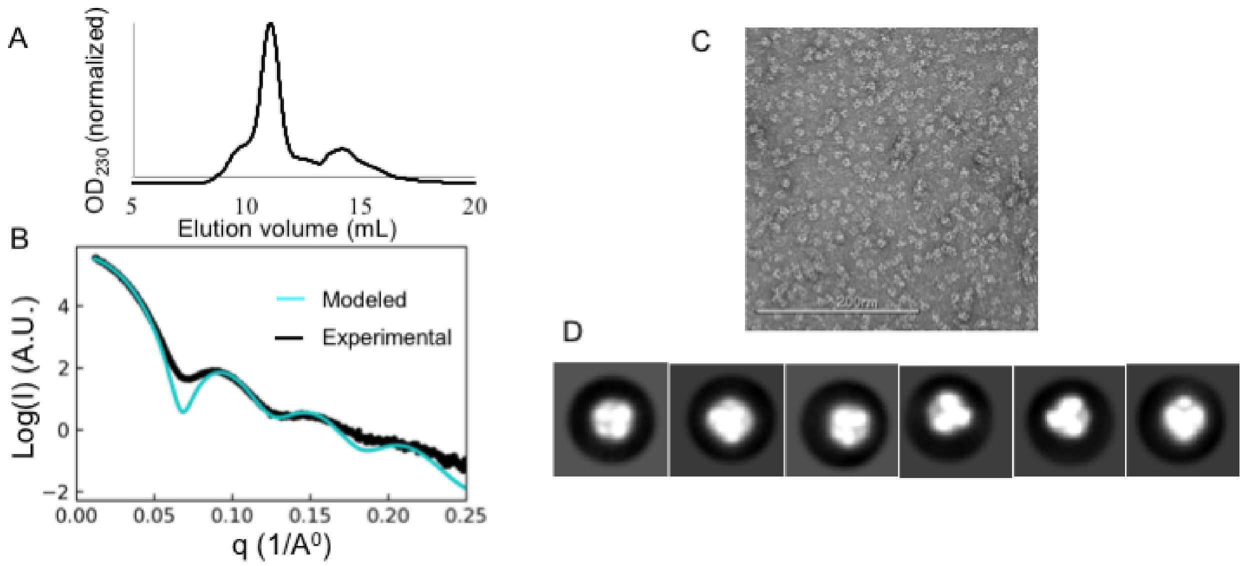

**Figure S20**: **SEC, SAXS, and Negative stain characterization of T_Wm-1606 tetrahedron**. A) SEC, B) SAXS, C) Representative micrograph, and D) 2D class averages.

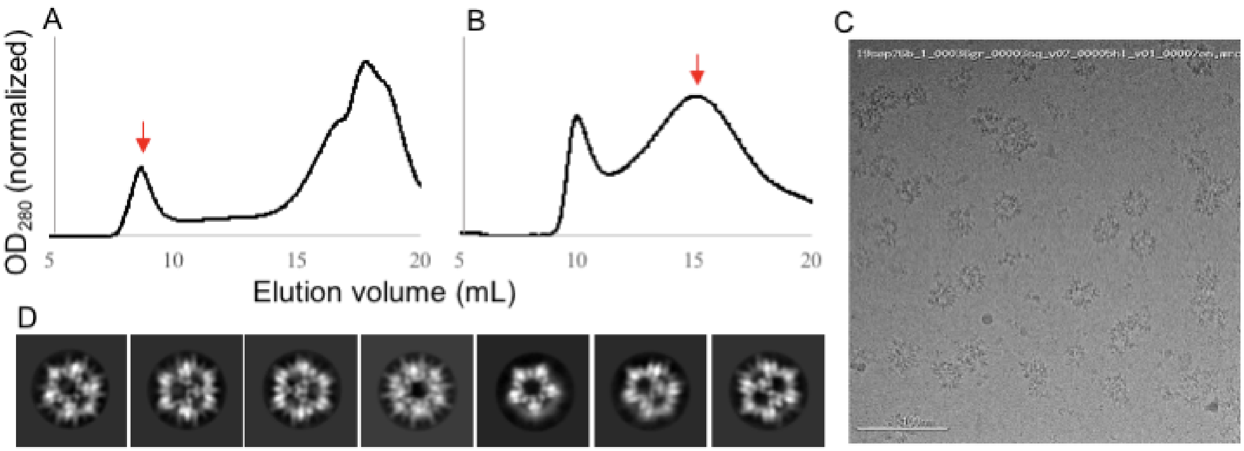

**Figure S21**: **SEC and Cryo electron microscopy characterization of I32_Wm-42 icosahedral nanocage**. SEC of I32_Wm-42 A) after Ni-NTA purification (Superose 6), and B) collected void fraction from A (red arrow) to re-run on Sephacryl 500. Fractions ~15mL were collected for further analysis (red arrow). C) Representative micrograph; D) class averages.

### 3) SUPPLEMENTARY TABLES

**Table S1:** Crystallographic Data Collection and Refinement Statistics

| | **C3_nat_HF-0005** (PDB: 6XH5) | **C3_HF_Wm-0024A** (PDB: 6XI6) | **C3_HD-1069** (PDB: 6XT4) | **C3_Crn-05** (PDB: 6XNS) |
|---|---|---|---|---|
| **Data collection** | | | | |
| Space group | $P4_32_12$ | *R3 :H* | *R3 :H* | $P22_12_1$ |
| Cell dimensions | | | | |
| *a*, *b*, *c* (Å) | 166.77, 166.77, 223.51 | 101.97, 101.97, 78.44 | 107.31, 107.31, 56.06 | 112.13, 145.25, 161.89 |
| *α*, *β*, *γ* (°) | 90, 90, 90 | 90, 90, 120 | 90, 90, 120 | 90, 90, 90 |
| Resolution (Å) | 78.12 - 3.32 (3.43 – 3.32)[a] | 38.48 - 2.69 (2.78 - 2.69) | 35.78 - 2.4 (2.486 - 2.4) | 46.39 - 3.19 (3.30 - 3.19) |
| No. of unique reflections | 47181 (4621) | 8434 (844) | 9405 (928) | 44729 (4436) |
| $R_{merge}$ | 0.238 (1.824) | 0.071 (0.577) | 0.139 (0.467) | 0.098 (2.98) |
| $R_{\text{pim}}$ | 0.046 (0.348) | 0.038 (0.324) | 0.06529 (0.216) | 0.035 (1.048) |
| $I/\sigma(I)$ | 17.98 (2.09) | 11.4 (2.3) | 6.13 (2.26) | 13.18 (0.93) |
| $CC_{1/2}$ | 0.986 (0.723) | 0.997 (0.889) | 0.993 (0.922) | 0.999 (0.247) |
| Completeness (%) | 99.88 (99.98) | 99.59 (99.41) | 99.27 (99.15) | 99.79 (99.57) |
| Redundancy | 27.2 (28.4) | 4.8 (4.8) | 5.6 (5.6) | 8.9 (9.0) |
| **Refinement** | | | | |
| Resolution (Å) | 78.12 – 3.32 | 38.48 - 2.69 | 35.78 - 2.4 | 46.39 - 3.19 |
| No. of reflections | 47141 | 8412 | 9338 | 44729 |
| $R_{\text{work}}$ / $R_{\text{free}}$ (%) | 22.17 / 26.48 (30.12 / 37.64) | 22.10 / 27.72 (33.81 / 36.32) | 22.75 / 27.30 (30.82 / 34.85) | 27.08 / 29.56 (41.15 / 40.11 ) |
| No. atoms | 15883 | 2056 | 1618 | 12559 |
| Protein | 15834 | 2043 | 1614 | 12559 |
| Water | 49 | 13 | 4 | 0 |
| Ramachandran Favored/allowed Outlier (%) | 95.47/4.29 00.24 | 98.50/1.50 00.00 | 99.56/0.44 00.00 | 98.77/1.23 00.00 |
| R.m.s. deviations | | | | |
| Bond lengths (Å) | 0.002 | 0.001 | 0.004 | 0.002 |
| Bond angles (°) | 0.470 | 0.330 | 0.56 | 0.47 |
| $B_{\text{factors}}$ (Å$^2$) | | | | |
| Protein | 117.47 | 74.96 | 54.58 | 130.31 |
| Water | 89.90 | 69.13 | 53.08 | - |

**Table S2:** CryoEM data collection parameters for C4_nat_HF-7900 and C5_HF-3921

| | **C4_nat_HF-7900** (6XSS, EMD-22305) | **C5_HF-3921** (EMD-22306) |
|---|---|---|
| Microscope | Talos Arctica | Titan Krios |
| Electron energy | 200 kV | 300 kV |
| Pixel size | 0.859 Å | 1.083 Å |
| Total electron dose | 57.05 $e^{-}/Å^{2}$ | 68.61 $e^{-}/Å^{2}$ |
| Number of frames in each movie | 40 | 50 |
| Exposure time | 2800 ms | 2500 ms |
| Defocus range | -0.2 - -4.2 μm | -1.9 - -5.0 μm |
| Tilt angle(s) | 0 ° | 0, 35 ° |
| Number of images acquired | 3,752 | 6,744 |
| Number of particles used in final map | 144,329 | 30,659 |
| Final map resolution (FSC = 0.143) | 3.70 | 8.06 |
| B-factor for map sharpening | -180 $Å^{2}$ | -500 $Å^{2}$ |
| Sphericity of 3DFSC | 0.895 | 0.786 |
| EMDB entry number (map) | EMD-22305 | EMD-22306 |
| EMPIAR entry number (data) | XXX | XXX |

**Table S3:** Model statistics for C4_nat_HF-7900 cryoEM structure

| | |
|---|---|
| Map CC (mask) | 0.78 |
| Map CC (volume) | 0.77 |
| Map CC (peaks) | 0.63 |
| rmsd (bonds) | 0.003 Å |
| rmsd (angles) | 0.605° |
| Ramachandran plot values | |
| outliers | 0.00% |
| allowed | 2.52% |
| favored | 97.48% |
| Rotamer outliers | 0.00% |
| C-beta deviations | 0.00% |
| Overall score (Molprobity[16]) | 2.04 |
| PDB ID | 6XSS |

### Design Construct Renaming

#### HelixDock

| Published name | Original Name | Published name | Original Name |
|---|---|---|---|
| C2_HD-1091 | YH_1BH-91 | C3_HD-1064 | YH_1BH-64 |
| C2_HD-1092 | YH_1BH-92 | C3_HD-1066 | YH_1BH-66 |
| C2_HD-1093 | YH_1BH-93 | C3_HD-1068 | YH_1BH-68 |
| C2_HD-1096 | YH_1BH-96 | C3_HD-1069 | YH_1BH-69 |
| C3_HD-1005 | YH_1BH-05 | C6_HD-1010 | YH_1BH-10 |
| C3_HD-2019 | UN_1BH-19 | C6_HD-1011 | YH_1BH-11 |
| C3_HD-1046 | YH_1BH-46 | C6_HD-1013 | YH_1BH-13 |
| C3_HD-1053 | YH_1BH-53 | C6_HD-3014 | C6-14 |
| C3_HD-1058 | YH_1BH-58 | C6_HD-3019 | C6-19 |

#### HelixFuse

| Published name | Original Name |
|---|---|
| C5_HF-2101 | C5-21-01 |
| C5_HF-3921 | C5-39-21 |
| C5_HF-0007 | C5_HFuse-0007 |
| C5_HF-0016 | C5_HFuse-0016 |
| C5_HF-0019 | C5_HFuse-0019 |
| C5_HF-0032 | C5_HFuse-0032 |
| C6_HF-0069 | C6-69 |
| C6_HF-0075 | C6-75 |
| C6_HF-0080 | C6-80 |
| C3_nat_HF-0005 | 1wa3_HFuse_BA-05 |
| C4_nat_HF-7900 | C4-79 |

#### Worms Designs

| Published name | Original Name |
|---|---|
| C3_Crn-05 | C3_hetC2_HFuse-05, C3_crown-05 |
| C5_Crn-07 | C5_hetC2_HFuse-07, C5_crown-07 |
| C5_Crn_HF-12 | crn_arm-12, C5_crown-07_HFuse-12 |
| C5_Crn_HF-26 | crn_arm-26, C5_crown-07_HFuse-26 |
| C5_Crn_HF-12_26 | crn_arm-12_26 |

| Published name | Original Name |
|---|---|
| D2_Wm-01 | D2-1 |
| D2_Wm-01_trunc | D2-1_trunc |
| D2_Wm-02 | D2-2 |
| T_Wm-1606 | T16.6 |
| I32_Wm-42 | w2c_DHRsp-42 |
| C3_HF_Wm-0024A | w2c_DHRsp-24A_capped |

### 4) SUPPLEMENTARY MAIN-TEXT PROTEIN SEQUENCES

*<u>Underline</u> denotes added linker, start codon, and his-tag residues used for Ni-NTA purification.

#### HelixDock

>C3_HD-1069
<u>MGHHHHHHGG</u>NDEKEKLKELLKRAEELAKSPDPEDLKEAVRLAEEVVRERPGSNLAKKALEIIL
RAAEELAKLPDPKALIAAVLAAIKVVREQPGSNLAKKALEIILRAAEELAKLPDPLALAAAVVA
ATIVVLTQPGSELAKKALEIIERAAEELKKSPDPLAQLLAIAAEALVIALKSSSEETIKEMVKL
TTLALLTSLLILILILLDLKEMLERLEKNPDKDVIVKVLKVIVKAIEASVLNQAISAINQILLA
LSD

#### HelixFuse

>C3_nat_HF-0005
<u>MGHHHHHHGGS</u>SEEEQERIRRILKEARKSGTEESLRQAIEDVAQLAKKSQDSEVLEEAIRVILR
IAKESGSEEALRQAIRAVAEIAKEAQDSEVLEEAIRVILRIAKESGSEEALRQALRAVAEIAEE
AKDERVRKEAVRVMLQIAKESGSKEAVKLAFEMILRVVRIIAVLRANSVEEAKEKALAVFEGGV
LAIEITFTVPDADTVIKELSFLEKEGAIIGAGTVTSVEQCRKAVESGALFIVSPHLDEEISQFC
DEAGVAYAPGVMTPTELVKAMKLGHRILKLFPGEVVGPQFVKAMKGPFPNVRFVPTGGVNLDNV
AEWFKAGVLAVGVGSALVKGTPDEVREKAKAFVEKIKAA

>C4_nat_HF-7900
MASSWVMLGLLLSLLNRLSLAAEAYKKAIELDPNDALAWLLLGSVLLLLGREEEAEEAARKAIE
LKPEMDSARRLEGIIELIRRAREAAERAQEAAERTGDPRVRELARELKRLAQEAAEEVRRDPDS
KDVNEALKLIVEAIEAAVRALEAAERTGDPEVRELARELVRLAVEAAEEVQRNPSSSDVNEALK
LIVEAIDAAVRALEAAEKTGDPEVRELARELVRLAVEAAEEVQRNPSSEEVNEALKDIVKAIQE
AVESLREAEESGDPEKREKARERVREAVERAEEVQRDPSS<u>GGSWGLEHHHHHH</u>

>C5_HF-3921
<u>MGHHHHHHGSGSENLYFQGGS</u>SDLQEVADRIVEQLKREGRSPEEARKEARRLIEEIKQSAGGDS
ELIEVAVRIVKELEEQGRSPSEAAKEAVELIERIRRAAGGDSELIEVAVRIVKELEEQGRSPSE
AAKEAVELIERIRRAAGGDSELIEVAVRIVKELEEQGRSPSEAAKEAVELIERIRRAAGGDSEL
IEVAVRIVKELEEQGRSPSEAAKEAVELIERIRRAAGGDSELIEVAVRIVKFLEEAGMSPSEAA
KVAVELIERIRRAAGGDSELIEKAVRIVRRLERRGLSPAEAAKIAVAIIAAEVLSREAEKIREE
TEEVKKEIEESKKRPQSESAKNLILIMQLLINQIRLLALQIQMLRLQLEL

>C5_HF-2101

MGHHHHHHGSGSENLYFQGGSSEKEKVEELAQRIREQLPDTELAREAQELADEARKSDDSEALKVVYLALRIVQQLPDTELAREALELAKEAVKSTDSEALKVVYLALRIVQQLPDTELAREALELAKEAVKSTDSEALKVVYLALRIVQQLPDTELAREALELAKEAVKSTDSEALKVVYLALRIVQQLPDTELAREALELAKEAVKSTDSEALKVVYLALRIVQQLPDTELAREALELAKEAVKSTDSEALKVVYLALRIVQLLPDTDLARKALELAKEAVKMDDQEVLKVVYKALQIVADKPNTEEADEALRDARLKLEAARLRREMEKIREETEEVKKEIEESKKRPQSESAKNLILIMQLLINQIRLLALQIRMLDLQLKL

>C5_HF-0019

MGHHHHHHGGSENLYFQSGGNDEKEKLKELLKRAEELAKSPDPEDLKEAVRLAEEVVRERPGSNLAKKALEIILRAAEELAKLPDPEALKEAVKAAEKVVREQPGSNLAKKAQEIILRAAEELAKLEDEEALKEAIKAAEKVIELEPGSELAKEAKRIIEKAAKMLADILRKEMEKIREETEEVKKEIEESKKRPQSESAKNLILIMQLLINQIRLLALQIRMLVLQLIL

>C6_HF-0075

MGHHHHHHGWSGSIQEKAKQSVIRKVKEEGGSEEEARERAKEVEERLKKEADDSTLVRAAAAVVLYVLEKGGSTEEAVQRAREVIERLKKEASDSTLVRAAAAVVLYVLEKGGSTEEAVQRAREVIERLKKEASDSTLVRAAAAVVLYVLEKGGSTEEAVQRAREVIERLKKEASDSTLVRAAAAVVLYVLEKGGSTEEAVQRAREVIERLKKEASDSTLVRAAAAVVLYVLEKGGSTEEAVQRAREVIERLKKEASDSTLVRAAAAVVLYVLEKGGSTEEAVDRAREVIEALKKFANDEEEIRRAAKVVLKVLETGGSVEEAMIRAALEILLDMLKEAAKKLKKLEDKTRRSEEISKTDDDPKAQSLQLIAESLMLIAESLLIIAISLLLSSLAG

>C6_HF-0080

MGHHHHHHGWSGSTKEKARQLAEEAKETAEKVGDPELIKLAEQASQEGDSEKAKAILLAAEAARVAKEVGDPELIKLALEAARRGDSEKAKAILLAAEAARVAKEVGDPELIKLALEAARRGDSEKAKAILLAAEAARVAKEVGDPELIKLALEAARRGDSEKAKAILLAAEAARVAKEVGDPELIKLALEAARRGDSEKAKAILLAAEAARVAKEVGDPELIKLALEAARRGDSEKAKAILLAAEAARVAKEAGIPEMIKAALRAARLGASDAAQAILEAADEARKAREEGDKKKEKSAELKALLALAKVKLKRLEDKTRRSEEISKTDDDPKAQSLQLIAESLMLIAESLLIIAISLLLSSDAG

**Crowns**

>C3_Crn-05

MGDRSDHAKKLKTFLENLRRHLDRLDKHIKQLRDILSENPEDERVKDVIDLSERSVRIVKTVIKIFEDSVRKLLKQINKEAEELAKSPDPEDLKRAVELAEAVVRADPGSNLSKKALEIILRAAAELAKLPDPDALAAAARAASKVQQEQPGSNLAKAAQEIMRQASRAAEEAARRAKETLEKAEKDGDPETALKAVETVVKVARALNQIATMAGSEEAQERAARVASEAARLAERVLELAEKQGDPEVARRAREL

QEKVLDILLDILEQILQTATKIIDDANKLLEKLRRSERKDPKVVETYVELLKRHERLVKQLLEI
AKAHAEAVEGGSLEHHHHHH

>C5_Crn-07
MGDRSEHAKKLKTFLENLRRHLDRLDKHIKQLRDILSENPEDERVKDVIDLSERSVRIVKTVIK
IFEDSVRKLEKQILKEAEELAKSPDPEDLKRAVELARAVIEANPGSNLSRKAMEIIERAARELS
KLPDPEAQRTAIEAASQLATMAAATGNTDQVRRAAELMKEIARLAGTEEAKDLALDALLDVLET
ALQIATKIIDDANKLLEKLRRSERKDPKVVETYVELLKRHEEAVRLLLEVAKTHADIVEGGSLE
HHHHHH

>C5_Crn_HF-12
MGDRSEHAKKLKTFLENLRRHLDRLDKHIKQLRDILSEHPHDERVKDVIDLSERSVRIVKKVIK
IFEDSVRELEKMILKEAEELAKSPDPEDLKRAVELARAVIEANPGSNLSRKAMEIIERAARELS
KLPDPEAQRTAIEAASQLATMAAATGNTDQVRRAAKLMMRIAILAGTEEASDLALDALLDVLET
ALQIATKIIDDANKLLEKLRRSHHHDPKVVETYVELLKRHEEAVRLLLDVAIMHALIVVMQDAI
EAAREGDKDRARKALQDALELARLAGTTEAVEAALLVVEAVAVAAARAGATDVVREALEVALEI
ARESGTTEAVKLALEVVASVAIEAARRGNTDAVREALEVALEIARESGTEEAVRLALEVVKRVS
DEAKKQGNEDAVKEAEEVRKKIEEES

>C5_Crn_HF-26
MGTESKVLEAEMSIKKAEWSAREGNPEKATEDLMRAMLLIRELDVLAQKTGSAEVLVKAAALAE
KLAKVAREVGDPEMAREAEKLARALAAKLLSMHAKLLATFLENLRRHLDRLDKHIKQLRDILSE
HPHDERVKDVIDLSERSVRIVKTVIKIFEDSVRKLLKEMLKRAEELAKSPDPLDLKAAVDVARA
VIEANPGSNLSRKAMEIIERAARELSKLPDPLAIATAIEAASQLATMAAATGNTDQVRRAAELM
KEIARLAGTDLAKAAALLALLRVLETALQIATKIIDDANKLLEKLRRSHHHDPKVVETYVELLK
RHEEAVRLLLEVAKTHADIVE

>C5_Crn_HF-12_26
MGTESKVLEAEMSIKKAEWSAREGNPEKATEDLMRAMLLIRELDVLAQKTGSAEVLVKAAALAE
KLAKVAREVGDPEMAREAEKLARALAAKLLSMHAKLLATFLENLRRHLDRLDKHIKQLRDILSE
HPHDERVKDVIDLSERSVRIVKKVIKIFEDSVRELLKMMLKRAEELAKSPDPEDLKAAVDVARA
VIEANPGSNLSRKAMEIIERAARELSKLPDPEAIATAIEAASQLATMAAATGNTDQVRRAAKLM
MRIAILAGTDLASAAALDALLRVLETALQIATKIIDDANKLLEKLRRSHHHDPKVVETYVELLK
RHEEAVRLLLDVAIMHALIVVMQDAIEAAREGDKDRARKALQDALELARLAGTTEAVEAALLVV
EAVAVAAARAGATDVVREALEVALEIARESGTTEAVKLALEVVASVAIEAARRGNTDAVREALE
VALEIARESGTEEAVRLALEVVKRVSDEAKKQGNEDAVKEAEEVRKKIEEES

**Dihedral rings**

>D2_Wm-01A
MGTREESLKEQLRSLREQAELAARLLRLLKELERLQREGSSDEDVRELLREIKELVAEIIKLIM
EQLLLIAEQLLGRSEAAELALRAIRLALELCRQSTDLEECLRLLKTAIKALENALRHPDSTTAK
ARLMAITARLLAQQLRTQHPDSQAARDAEKLADQAERAVRLATRLYEEHPNAEISEMCSQAAYA
AALMASIAAILAQRHPDSQIARDLIRLASELAEMVKRMCERGGSWGLEHHHHHH

>D2_Wm-01B
MGTREELAKELLRSLREQAESLARQLRLLKELERLQREGSSDEDVRELLREIKELAAEQIKLIM
EQLLLIAELTLGRSEAAELALDAIRQALEACRTMDNQEACTRLLKLAIQMLELATRAPDAEAAK
LALEAAKKAIELANRHPGSQAAEDATKLAQQAMEAVRLALKLYEEHPNADIADLCRRAAAEAAE
AASKAAELAQRHPDSQAARDAIKLASQAAEAVKLACELAQEHPNADKAKLCILLASAAALLASI
AAMLAQRHPDSQEARDMIRIASELAELVKEICER

>D2_Wm-02A
MGTREEIIRELARSLAEQAELTARLERSLREQERLQREGSSDEDVRELIREQKELVREILKLIA
EQILLIAELLLASTRSEAAELALRAIRNAIEACKNADNEEMCRQLMRMAQNALELATQAPDAEA
AKAALRAIDLAVELASRHPGSQAADDALKLAQQAAEAVKLALDLYREHPNADIADLCRKAAKEA
AEAASKAAELAQRHPDSQAARDAIKLASQAAEAVKLACELAQEHPNAEIAKMCILAASAAALMA
SIAAILAQRHPDSQIARDLIRLASELAEMVKRMCERGGSWGLEHHHHHH

>D2_Wm-02B
MGTREELAKELLRSLREQAESLARQLRLLKELERLQREGSSDEDVRELLREIKELAAEQIKLIM
EQLLLIAELMLGRSEAAELALEAIRLALELCRQSTDQEQCTDLLRQATEALETATRYPDDTNAK
AKLMAITARLLAQQLRTQHPDSQAARDAEKLADQAEKAVRLAKRLYEEHPNADKSELCSQLAYA
AALLASIAAMLAQRHPDSQEARDMIRIASELAELVKEICER

>D2_Wm-01_truncA
MGTREESLKEQLRSLREQAELAARLLRLQREGSSDEDVKELVAEIIKLIMEQLLLIAEQLLGRS
EAAELALRAIRLALELCRQSTDLEECLRLLKTAIKALENALRHPDSTTAKARLMAITARLLAQQ
LRTQHPDSQAARDAEKLADQAERAVRLATRLYEEHPNAEISEMCSQAAYAAALMASIAAILAQR
HPDSQIARDLIRLASELAEMVKRMCERGGSWGLEHHHHHH

>D2_Wm-01_truncB
MGTREELAKELLRSLREQAESLARQLRLQREGSSDEDVKELAAEQIKLIMEQLLLIAELTLGRS
EAAELALDAIRQALEACRTMDNQEACTRLLKLAIQMLELATRAPDAEAAKLALEAAKKAIELAN
RHPGSQAAEDATKLAQQAMEAVRLALKLYEEHPNADIADLCRRAAAEAAEAASKAAELAQRHPD
SQAARDAIKLASQAAEAVKLACELAQEHPNADKAKLCILLASAAALLASIAAMLAQRHPDSQEA
RDMIRIASELAELVKEICER

**Point Group nanocage**

>T_Wm-1606
MGDEEKKKELLKQLEDSLIELIRILAELKEMLERLEKNPDKDTIVKVLKVIVKAIEASVANQAI
SAMNQGADANAKDSDGRTPLHHAAEAGAAAVVKVAIDAGADVNEKDSDGRTPLHHAAENGHAEV
VTLLIEKGADVNEKDSDGRTPLHHAAENGHDEVVLILLLKGADVNAKDSDGRTPLHHAAENGHK
RVVLVLILAGADVNTSDSDGRTPLDLAREHGNEEVVKALEKQGGWLEHHHHHH

>I32_Wm-42A
MGGSELEIVIRLQILNLELARKLLEAVARLQELNIDLVRKTSELTDEKTIREEIRKVKEESKRI
VKEAEDEIKKAALISADLAAKAIKRAIDRAKKLLEKGEKEDAEDVLREARSAIRLVTELLERIA
KNSSTPEEALRAAELLVRLIILLIKIAALLAAAGNKEEADKVLDEAKELIERVRELLEKISKNS
DTPELSKRAKELELILRLADLAIKAMKNTGSDEARQAVKEMARLAKEALEMGM**S**EAAKAAIELL
ELLAEAFAGSDVASLAVKAIAKIAETALRNG**S**

*bolded Ser residue denotes additional mutation of Cys to remove a disulfide bond at the interface termini

>I32_Wm-42B
MG**S**DTAKEAIQRLEDLARKYSGSDVASLAVKAIEKIARTAVENG**S**EETAEEAEKRLRELAEDYQ
GSNVASLAASAIAEIAAARARFAAREMGDPRVEEIAKELERLAKEAAERVERRPDSEEDYRKLE
LAALIIKLFVSLLKQKRLAERLKELLRELERLQREGSSDEDVRELLREIKELVEEIEKLARKQE
YLVTELAKMMGGSGGSGGSGGSLEHHHHHH

*bolded Ser residue denotes additional mutation of Cys to remove a disulfide bond at the interface termini

>C3_HF_Wm_0024A
MGKELEIVARLQQLNIELARKLLEAVARLQELNIDLVRKTSELTDEKTIREEIRKVKEESKRIV
EEAEQEIRKAEAESLRLTAEAAADAARKAALRMGDERVRRLAAELVRLAQEAAEEATRDPNSSD
QNEALRLIILAIEAAVRALDKAIEKGDPEDRERAREMVRAAVRAAELVQRYPSASAANEALKAL
VAAIDEGDKDAARCAEELVEQAEEALRKKNPEEARAVYEAARDVLEALQRLEEAKRRGDEEERR
EAEERLRQACERARKKNGGSLEHHHHHH
